# RNA secondary structures are conserved but random

**DOI:** 10.1101/2025.08.18.670923

**Authors:** Samuel H. A. von der Dunk, Nora S. Martin, Kamal Dingle, Ard A. Louis

## Abstract

Noncoding RNAs perform a wide range of essential biological functions, and their secondary structures are often conserved by purifying selection. However, such conservation does not necessarily imply that positive selection shaped their evolutionary origins. Here, we test for global signs of positive selection by studying the distribution of secondary structures in naturally occurring noncoding RNA. We find that, to first order, these structures are statistically indistinguishable from those produced by a relatively small set of randomly generated sequences. The distributions are, however, profoundly shaped by a strong bias in the arrival of phenotypic variation, such that only an exponentially small subset of all possible structures is likely to occur in nature. In other words, the secondary structure repertoire of natural noncoding RNAs largely reflects this developmental bias rather than further adaptive fine-tuning. Detecting genuine signatures of selection, beyond randomness, in the distribution of secondary structures, therefore requires careful calibration against appropriate null models that account for the underlying bias. We perform a large-scale and detailed analysis of four extensive datasets covering a wide spectrum of functional RNA classes. We describe one potential signature of adaptation on structure: archaeal ribosomal RNA structures are simpler and more robust than predictions of the sequence null model, and hyperthermophiles are less complex than archaea in other niches, but the effects are relatively small. This example illustrates the difficulty of inferring a creative role for natural selection in shaping evolutionary outcomes for RNA secondary structure.

## Introduction

The molecular revolution in biology was driven by the principle that biomolecular function is mediated by structure. Indeed, some of the most significant advances in molecular biology arose from solving three-dimensional structures, revealing critical insights into their functions. This tight coupling between structure and function helps explain why protein and RNA structures are frequently conserved over evolutionary timescales, even when their primary sequences are not (Graur and Li, 2000).

A foundational principle of Darwinian evolution is that biological function emerges through adaptive processes that incrementally enhance fitness. Well-established methods exist to detect positive selection in sequences that encode proteins (Graur and Li, 2000). At a more fine-grained level, empirical fitness landscapes can be directly measured (De Visser and Krug, 2014). The intimate relationship between structure and function suggests that molecular structures have also undergone adaptive refinement. Indeed, evidence for selection on structure has been demonstrated, for example, through ancestral protein reconstruction (Ortlund et al., 2007), directed evolution (Romero and Arnold, 2009), and comparative genomics (e.g. Szilágyi and Závodszky, 2000; Deng et al., 2010). However, it is also well recognized that neutral processes, particularly in eukaryotes with relatively small effective population sizes, can produce molecular function and structural complexity in the absence of positive selection (Lynch, 2007; Fernández and Lynch, 2011; Lukeš et al., 2011; Brunet et al., 2021; Torri et al., 2022).

Here we focus on the structure of noncoding RNA (ncRNA), functional molecules which, like proteins, are essential for many crucial cellular processes (Geisler and Coller, 2013; Santosh et al., 2015; Mattick et al., 2023). The capacity of RNA to both perform catalytic functions and encode information, as well as its central role in translation, provides indirect evidence for the existence of an RNA world (Gilbert, 1986), a prebiotic era before the advent of proteins in which biology consisted almost exclusively of RNA. The functional importance of ncRNA is supported in many cases by conservation of secondary structure (Smith et al., 2013; Rivas, 2021; Walter Costa, 2023; Mattick et al., 2023; Backofen et al., 2024), as in the famous clover-leaf shape of tRNA (Westhof et al., 2022).

Techniques for identifying positive selection in noncoding RNAs (ncRNAs) remain far less developed compared to methods used for proteins, and examples of adaptive evolution shaping RNA secondary structures are relatively rare (Walter Costa, 2023; Backofen et al., 2024). Instead of focusing on individual molecules, as is traditionally done, we draw inspiration from evolutionary developmental biology (evo-devo), where major insights have emerged by comparing observed biological forms to the full space of theoretically possible morphologies (Erwin, 2007). Such global comparisons can help distinguish the effects of developmental bias, reflecting biophysical constraints, from those of natural selection, reflecting the shaping role of the environment.

We extend this framework to RNA secondary structure, where fast and computationally efficient folding algorithms (Lorenz et al., 2011) make it possible to characterise the distribution of structural features across the entire universe of possible RNA folds. This tractability enables the construction of null models that explicitly exclude selection but account for biases inherent to RNA folding, which can be seen as a stripped-down version of the broader concept of developmental bias. The hypothesis we test is that signatures of adaptation can be detected by measuring deviations in the global structural repertoire of naturally occurring ncRNA relative to null expectations derived from models that omit natural selection. Given that RNA is thought to be highly evolvable (Schuster et al., 1994; Huynen, 1996; Schultes and Bartel, 2000; Greenbury et al., 2022), it would be surprising if extensive evolutionary histories in diverse environments would leave no detectable imprint on the structural diversity of natural ncRNA.

Previous work has already found remarkable similarities between the secondary structures of natural ncRNA and those produced by randomly generated RNA sequences (Fontana et al., 1993; Rivas, 2021; Smit et al., 2006; Jörg et al., 2008; Stich et al., 2008). In particular, natural ncRNA molecules fold into secondary structures that are exceptionally frequent in sequence space and are therefore readily found by random sampling of sequences (Jörg et al., 2008; Stich et al., 2008; Cowperthwaite et al., 2008; Dingle et al., 2015, 2022; Ghaddar and Dingle, 2023). Yet, with some exceptions, only relatively small ncRNA molecules with narrow functional and taxonomic coverage were considered in these studies. Here we present a comprehensive analysis for a much larger range of ncRNA lengths and classes, enabling us to examine the impact of natural selection on RNA secondary structure as a whole.

To characterise the universe of RNA structures we employ several metrics, placing particular emphasis on the distribution of structural complexity within RNA secondary structures, quantified by the compressibility of their dot-bracket representations. There are two main motivations for studying this measure.

The first is theoretical: recent work predicts that many genotype-phenotype maps exhibit a strong intrinsic bias towards simple phenotypes with low descriptional complexity (Dingle et al., 2018; Johnston et al., 2022). For RNA, this implies that secondary structures with highly compressible dot-bracket representations should appear with relatively high frequency when sequences are randomly sampled. However, this bias towards simplicity for each individual structure is countered by a second effect: there are many more complex structures than simple ones. The interplay between these two opposing forces— one favoring simplicity, the other favoring complexity—remains poorly understood and presents a rich avenue for investigation.

The second motivation is practical: our complexity measure offers a concise, coarse-grained summary of key structural features such as stacks and loops. By comparing complexity distributions of structures produced by a null model from randomly sampled RNA sequences to those of naturally occurring ncRNA, we can quantitatively evaluate the relative contributions of developmental bias and natural selection in shaping RNA secondary structure. Our main finding is both striking and thought-provoking: while developmental bias exerts a profound influence on evolutionary outcomes, we find remarkably little evidence for a constructive role of selection in shaping secondary structures across diverse classes of functional RNA.

## Results

### Natural RNA molecules have similar structures to those generated by random sequences

To explore the universe of RNA secondary structure folds, we develop two null models. For the *random phenotype null model*, we uniformly sample phenotypes from the set of all possible secondary structures using a technique from Burghardt and Hartmann (2007) (see Methods). For the *random sequence null model*, we generate random nucleotide sequences which are then folded into their minimum-free-energy secondary structure using the ViennaRNA package (Lorenz et al., 2011). The sequence null model is simple, yet has been shown to capture the spectrum of phenotypic variation produced by a population undergoing neutral evolution for RNA secondary structures (Schaper and Louis, 2014; Johnston et al., 2022) (see also Section F in the Supporting Information (SI)).

We compare the complexity distributions of natural ncRNA data to the two null models using our structural complexity measure *K*_struc_, which is calculated with Lempel-Ziv compression on the dot-bracket notation, following Dingle et al. (2018) (Methods). Initially, we focus on sequences of length *L* = 300, which is centrally positioned in the intermediate ncRNA length regime defined by the consensus statement in Mattick et al. (2023) and encompasses a wide range of functions for which structure is largely conserved. Later, we extend our analysis to the full RNA length range.

Natural ncRNA sequences were obtained from RNAcentral (The RNAcentral Consortium, 2021), filtered, separated into bacterial (*N* = 292) and eukaryotic sequences (*N* = 5159; Methods and Fig. S10), and folded into minimum-free-energy secondary structures (Lorenz et al., 2011). We find that the structural complexities of natural ncRNA molecules are remarkably similar to those produced by a sample of *N* = 10^5^ structures from the sequence null model (Fig. 1a–b). They are significantly simpler than the full range of possible structures represented by the phenotype null model. The complexity distribution of phenotypes that evolution has produced can be almost entirely explained by a strong developmental bias in the genotype-phenotype map.

**Figure 1.**
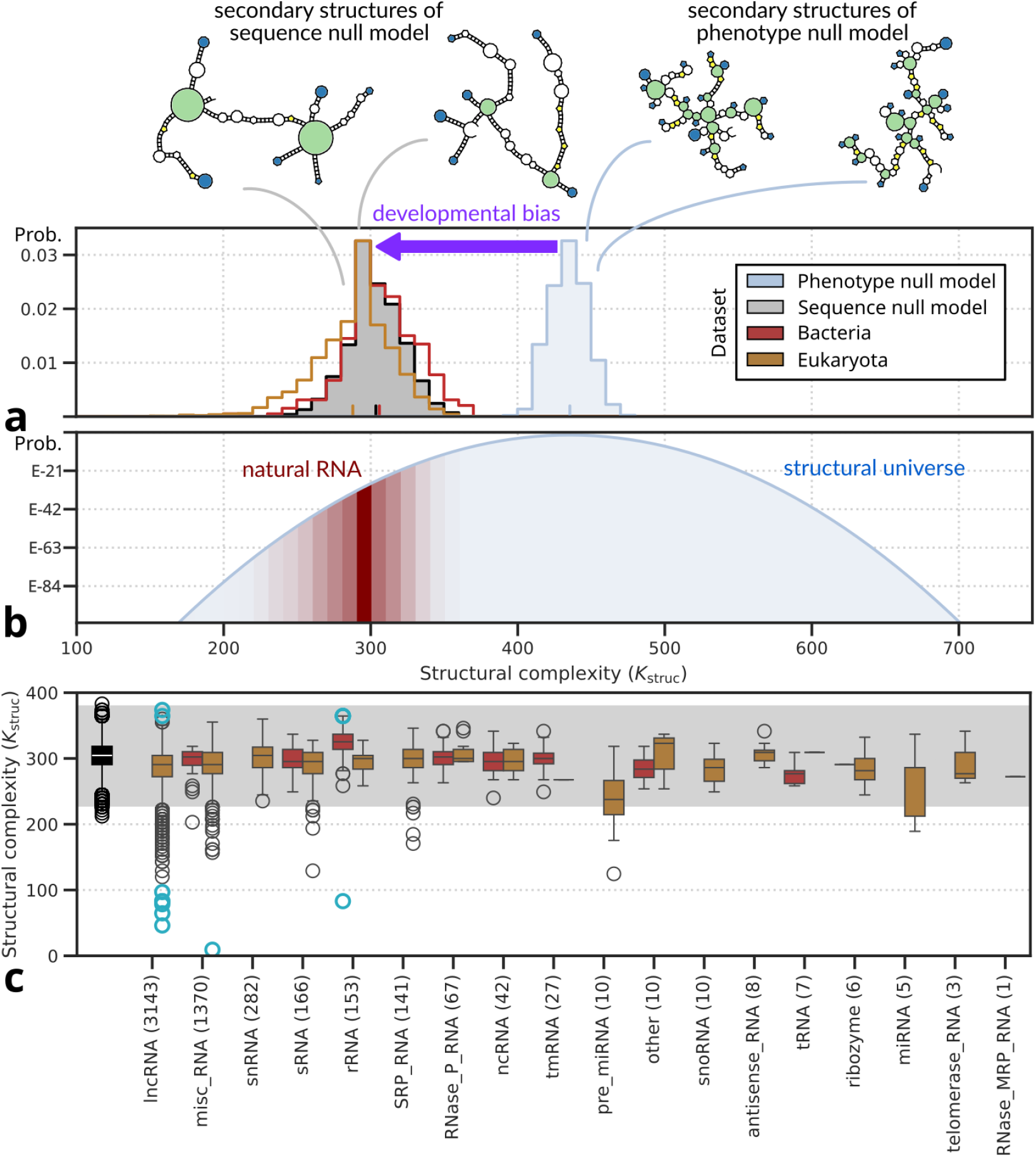
Probability distributions of structural complexity for natural RNA of length *L* = 300 are compared to the structural universe of RNA (phenotype null model) and phenotypes produced by random sequences (sequence null model) on (**a**) linear and (**b**) logarithmic scales. The impact of developmental bias (the difference between the two null models) is represented by a purple arrow. In panel (**b**), a normal distribution was fitted to extend the distribution produced by the phenotype null model. Natural RNA secondary structures of length *L* = 300 from RNAcentral closely resemble the structures produced by random sequences, but are drawn from an extreme tail of the set of all possible RNA structures (phenotype null model). Note that we have not yet taken into account biases in sequence composition which largely explain the small differences between bacteria and eukaryota (see Fig. 2). (**c**) Structural complexity distributions of different RNA types. The grey area marks the range of structural complexities found within four standard deviations of the mean for structures generated from the sequence null model (black in left-most box); cyan circles are the extremes of natural RNA (presented in Table S1 and Fig. S11). Boxes display the inter-quartile range and whiskers extend up to 1.5 times the length of the inter-quartile range.

The observed distribution arises from two competing effects (Johnston et al., 2022): complex structures, which form the majority of potentially realisable phenotypes, are rare in sequence space and hard to find when sampling random sequences. Conversely, simple structures are individually frequent in sequence space, but there are fewer such phenotypes. Thus, the average structural complexity of structures folded from random sequences is at an intermediate value (*K*_struc_ *≈* 300), but much lower than the mean (or mode) of all structures (*K*_struc_ *≈* 450); see SI Section A for more details.

The scale of the shift in structural complexity going from the sequence null model to the phenotype null model is worth emphasising. We find that the structural complexity of uniformly sampled phenotypes is well approximated by a normal distribution (see SI Section C) and that the average complexity of natural RNA is about 12.0 standard deviations lower than the average of the full universe of shapes. In other words, natural RNA are drawn from an extreme low-complexity tail of the full set of possible folds. To encounter a structure with complexity as low as that of a typical natural RNA molecule, we would have to sample on the order of 10^33^ random structures in the phenotype null model. Conversely, the strong bias of the sequence null model implies that the vast majority of theoretically possible secondary structures are unlikely ever to be found without directed selection.

Although most shapes from the universe of secondary structures are rare in sequence space, RNA is highly evolvable and could in principle navigate the sequence-structure map to find rare structures (Schuster et al., 1994; Huynen, 1996; Schultes and Bartel, 2000; Greenbury et al., 2022; Srivastava et al., 2025). Indeed, as we show in SI Section F, high- and low-complexity structures readily evolve under selection. Nevertheless, we find that the structural complexity of most natural RNA molecules lies within four standard deviations of the mean for structures generated from the sequence null model, and most of this range is covered with a sample of only 10^5^ molecules. Apparently, nature did not need to exploit the evolvability of RNA much. Instead, our observations imply that the functions performed by natural ncRNA are readily found by starting from random RNA sequences and as a consequence, natural ncRNA structures have remained within this set.

In our natural *L* = 300 RNA dataset, we encounter a range of different functional types, although the coverage is heterogeneous (Fig. 1c). To first order, the complexities of all the different types are quite similar to one another. For those types which are found in both bacteria and eukaryotes, structural complexities are also typically similar between the two domains. Individual RNA molecules can have extreme structural complexities, as seen in eukaryotic long noncoding RNA (lncRNA) for instance. These deviations from the structural complexity distribution of structures generated from random sequences could indicate that selection has played a role in finding such special structures. However, as we shall see below, many of these outliers are characterized by highly repetitive sequences, implying a role for biased mutational processes in their generation, rather than direct selection on structure (see also Table S1).

Although our structural complexity provides a single theoretically grounded measure for comparison between null models, it is also interesting to study other properties of RNA secondary structure. First, we show that mutational robustness, measured as the fraction of single-nucleotide mutants with the same structure as the original molecule, is similarly distributed for natural ncRNA and structures generated from the sequence null model (Fig. 2a), as observed for shorter RNA lengths (Dingle et al., 2015; Johnston et al., 2022, see also Fig. S13). Further, the abundance profiles of several secondary structure motifs of natural RNA are remarkably similar to those generated by the sequence null model, in line with their closely aligned structural complexity distributions (Fig. 1). Moreover, by comparing the null models (Fig. 2b–f; cf. Stich et al., 2008; Dingle et al., 2015; Ghaddar and Dingle, 2023), we pinpoint the effects of developmental bias for each of these structural motifs.

**Figure 2.**
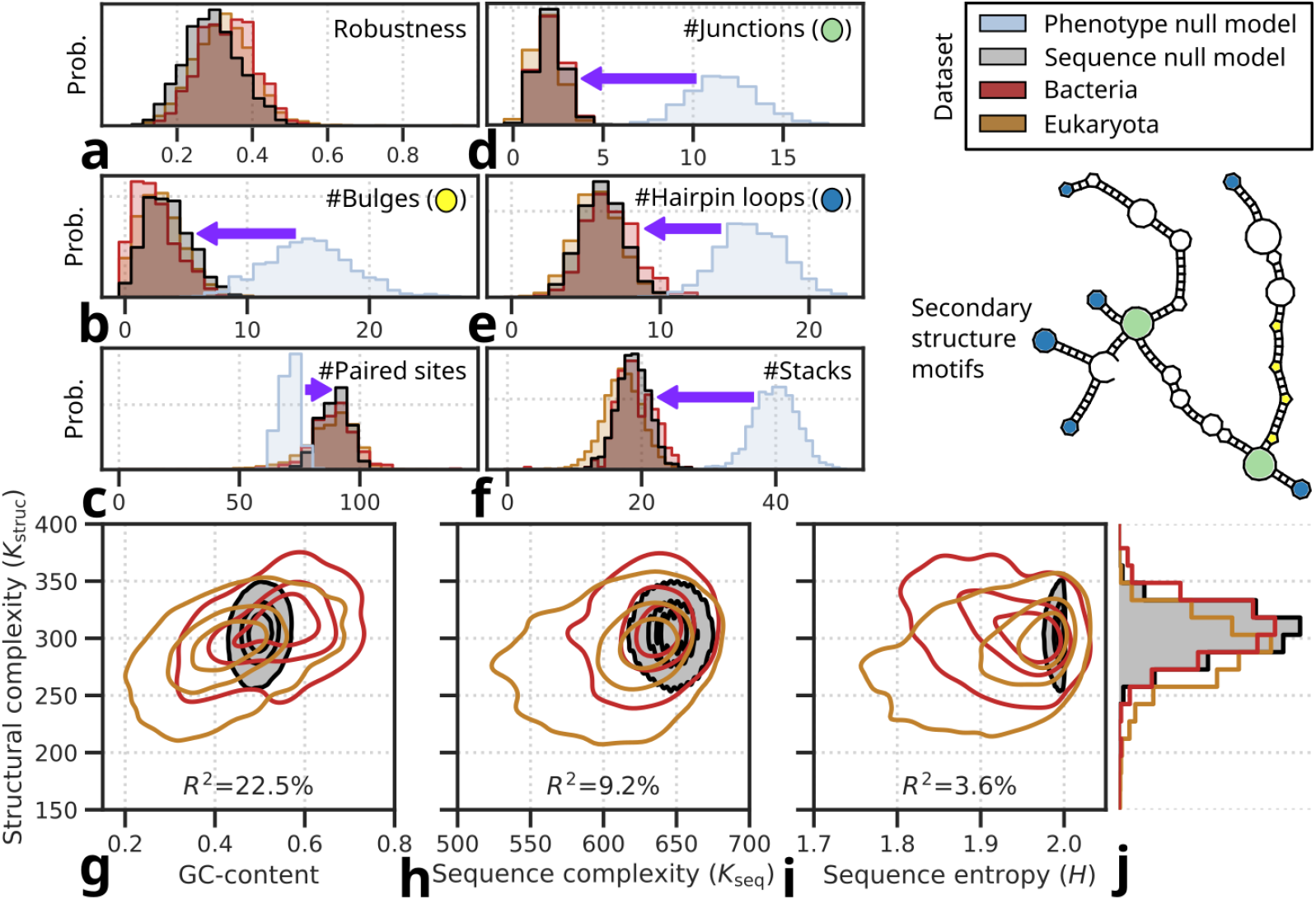
Characterization of mutational robustness, structural motifs and sequence features in random and natural RNA of length *L* = 300. (**a**) Probability distribution of mutational robustness (fraction of mutants with identical secondary structure) for the three sequence sets. (**b**–**f**) Probability distributions of secondary structure motif counts. Purple arrows represent developmental bias as in Fig. 1. (**g**–**j**) Kernel density plots showing correlations of structural complexity with three features of sequence composition, i.e. (**g**) GC-content, (**h**) sequence complexity, and (**i**) sequence entropy. Together, these features account for 27.3% of the total variation in structural complexity; similarly, *R*^2^ values represent variation explained across all three RNA sequence sets. For structural properties (panels **a**–**f**), we analyzed sub-samples of *N* = 1000 for structures generated from both null models. Panels **g**–**j** display the same data as in Fig. 1 (i.e. *N* = 10^5^ for structures generated from the sequence null model), but for calculating R-squared, all three sequence datasets were down-sampled to the same size (*N* = 228); the number of bacterial RNA genes for which we were able to obtain genomic GC-content; see Fig. S12). See Fig. S13 for the same analysis for RNAs of length *L* = 100.

### The effect of sequence composition on structure distributions

Despite impressive similarities between the structures of bacterial and eukaryotic ncRNA and structures generated from random sequences, subtle differences remain in their structural complexity distributions (Fig. 1a, 2j). We find that these differences in structural complexity are largely accounted for by biases in underlying sequence composition. Specifically, structural complexity correlates with GC-content, sequence complexity, and sequence entropy (Fig. 2g–i). After correcting for these features of sequence composition, almost no differences remain in structural complexity distributions between structures generated from random sequences and bacterial and eukaryotic RNA (Table S2).

The correlation between GC-content and structural complexity follows from RNA folding physics since GC pairs form stronger bonds than AU pairs, and so sequences with more G’s and C’s are typically able to form more complex bonding patterns (Stich et al., 2008; Dingle et al., 2015). In our data, eukaryotic RNA genes feature lower GC-content than structures generated from random sequences leading to simpler structures while bacterial RNA genes feature higher GC-content than structures generated from random sequences leading to slightly more complex structures (Fig. 2g). As previously described in the literature, GC-content of RNA genes correlates with the GC-content of genomes (Muto and Osawa, 1987, Fig. S12), suggesting that some of the differences in RNA sequence composition may be driven by genome-wide GC biases (Romiguier and Roux, 2017). The GC-content of eukaryotic lncRNA in particular, follows genomic GC-content. For many other RNA types, GC-content deviates from the genomic context, and part of the reason for this is that selective constraints preserve secondary structure even when the background genomic GC-content changes (Smit et al., 2006; Stich et al., 2008).

While higher GC-content generally increases structural complexity, a sequence consisting entirely of G or C would not form any base pairs, resulting in the simplest possible secondary structure. This explains why larger sequence entropy (i.e. more balanced nucleotide fractions) allowing all three base pairs including the GU wobble pair to occur, promotes structural complexity. Our compressibility calculation can also be used directly on the sequence to measure the sequence complexity *K*_seq_ (see SI section B). Even with high entropy, sequences that are repetitive corresponding to low *K*_seq_ tend to produce low-complexity structures, as seen in the simplest natural RNA molecules in Fig. 1c (see also Fig. S11,

Table S1). For instance, the simplest sequence (sRNA) comes from the apicomplexan *Eimeria necatrix* and consists for 96% of repetitions of the pattern GCAGCT[AG]C[TC]. The RNA molecule folds onto itself, forming a mostly double-stranded RNA molecule with low structural complexity. Similarly, the simplest RNA structure (misc RNA) from the butterfly *Melitaea cinxia* has a very low sequence complexity, with 54 repetitions of the pattern [CU]AGA[AG] interspersed 3 times by the pattern UAAGAA for 96% of the sequence.

### Comparing RNA structure distributions for a wide range of lengths

To generalize our findings to lengths other than *L* = 300, we analyzed structures generated for a large range of sequence lengths, from *L* = 30 to *L* = 10, 000. While ViennaRNA is most accurate for shorter RNA sequences, we do not expect our results to be significantly impacted by technical limitations (see SI Section D for discussion, including validation with experimental structural data). We derived a scaling law between structural complexity and length and between sequence complexity and length (Fig. S14– 16). This scaling relationship was used to quantify the *relative* structural complexity 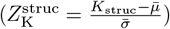 of an RNA molecule with complexity *K*_struc_ as the number of standard deviations *σ* above or below the mean 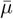 generated by structures of the same length produced by the sequence null model. We also generated random phenotype sets with the phenotype null model for lengths *L* = 30 to *L* = 500, which, due to computational constraints, was the upper limit for the algorithm of Burghardt and Hartmann (2007). With these methods, we can compare the relative structural complexity 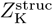 between RNA molecules of different lengths, as shown in Fig. 3.

**Figure 3.**
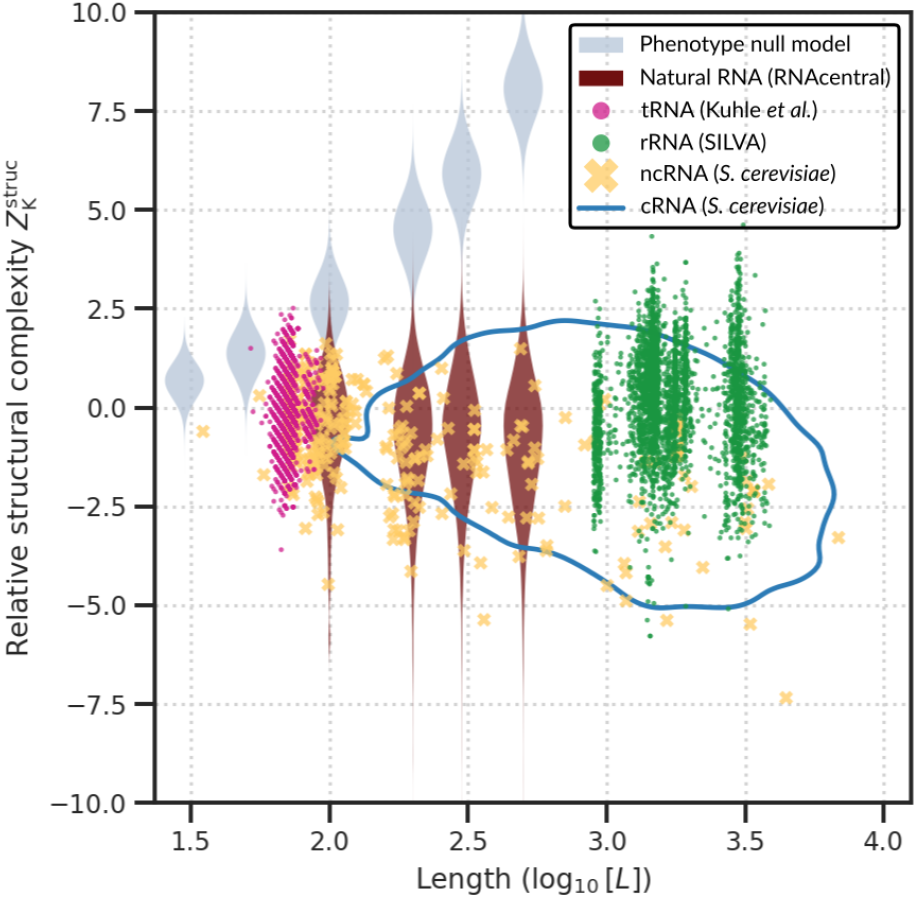
Structural complexity of natural RNA molecules is close to random expectation for many lengths. We compare the relative complexity 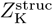 (the number of standard variations above or below the complexity of structures of the same length generated with the sequence null model) as a function of sequence length. Light grey violins show the distributions from the phenotype null model (*N* = 10^4^ each). The blue outline indicates the area where most coding RNA of *Saccharomyces cerevisiae* is found (*N* = 5511). Markers show length and complexity of individual RNA molecules from all three domains of life: noncoding RNA from RNA Central (*N* = 20940, maroon), of *S. cerevisiae* (*N* = 344, yellow) from SGD (Engel et al., 2022), as well as 20 tRNA types (*N* = 5992, pink) from (Kuhle et al., 2020), and 2 rRNA types (*N* = 3322, pink) from the SILVA dataset (Quast et al., 2012). All data is available online as `RNA complexity data.dat` (see Data Availability).

For all lengths, the structural complexities of phenotypes from the phenotype null model are clearly much larger than those generated by the sequence null model, as seen previously for *L* = 300. We also compared a diverse set of ncRNAs from the RNAcentral database (The RNAcentral Consortium, 2021), *S. cerevisiae* coding and ncRNA from the Saccharomyces Genome Database (SGD) (Engel et al., 2022), tRNA from (Kuhle et al., 2020), and rRNA from the SILVA database (Quast et al., 2012). Most RNA molecules have a structural complexity that lies within four standard deviations of the mean for structures generated by random sequences. Interestingly, some natural RNA molecules, i.e. those from RNAcentral and many long sequences from *S. cerevisiae*, exhibit a slight bias towards lower structural complexity, which is caused by low GC-content (see Fig. 4).

**Figure 4.**
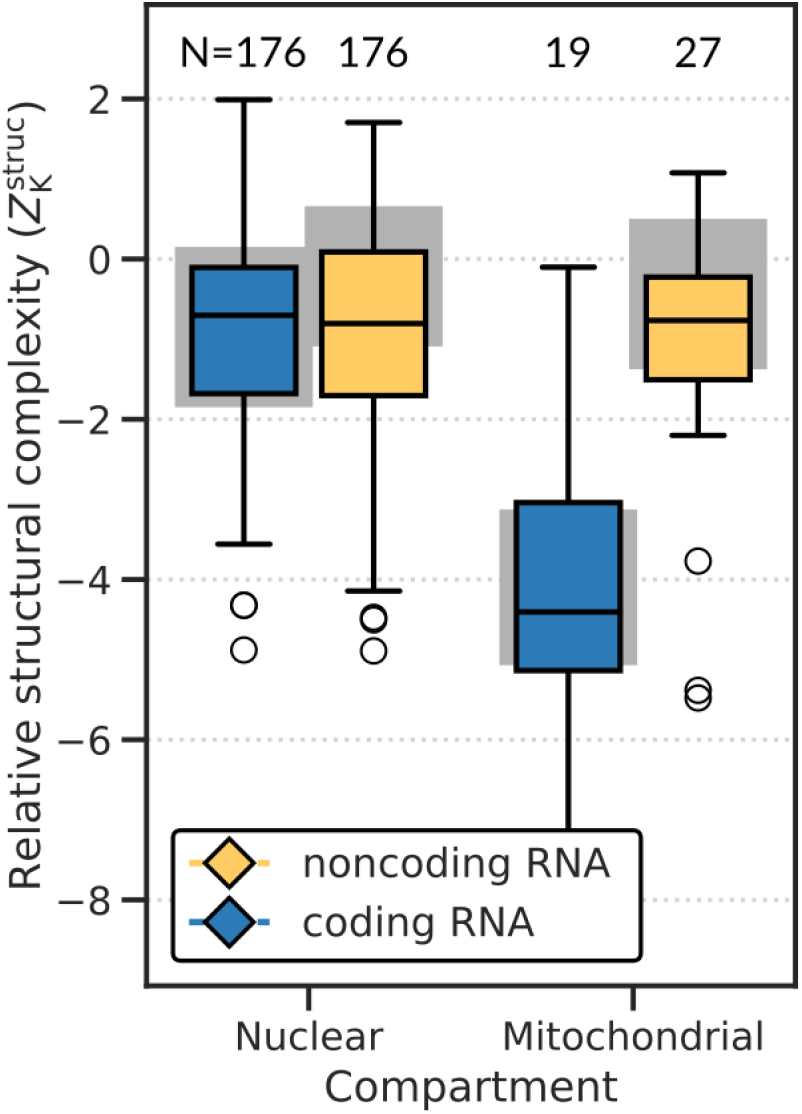
Mitochondrial noncoding RNA molecules for *S. cerevisiae* have maintained similar structural complexity as RNA molecules encoded on the nuclear genome. This is surprising as the mitochondrial genome is GC-poor and therefore prone to create RNA molecules with simple secondary structures, as seen in mitochondrial coding RNA. Colored boxes plus whiskers and outliers represent the relative structural complexity of RNA genes; grey boxes represent the relative structural complexity after shuffling the sequences, showing that sequence composition is sufficient to explain the complexity distributions of the natural genes. The three outliers of mitochondrial noncoding RNA are, from low to high complexity: 21S rRNA, 15S rRNA and RNase P; the remaining 24 genes are tRNA (see also Fig. S18).

A user-friendly online notebook (see Code Availability) allows users to compute the relative complexity (*Z*_K_) of the structure and sequence of any RNA molecule. These metrics help identify deviations from structural or sequence-level randomness, allowing users to assess whether a given RNA molecule exhibits unusually high or low complexity.

### Conservation of mitochondrial ncRNA structures in yeast

We next compare nuclear and mitochondrial coding and noncoding RNA in *S. cerevisiae*. We sampled from the full set of coding RNA (*N* = 5511 genes) a similar number to the set of ncRNA (*N* = 203, the number of genes for which we obtained genomic annotation), selecting for even representation of nuclear and mitochondrial genes and similar length distributions. We show a comparison between nuclear and mitochondrial RNA in Fig. 4. For nuclear RNA there are only modest deviations from the expectation of sequence null model, caused by the lower GC-content (about 40%, Fig. 4).

By contrast, mitochondrial coding RNA stands out from other RNA by its low structural complexity which we hypothesize is due to the low GC-content of the mitochondrial genome (about 25%, Fig. 4). Shuffling the mitochondrial coding RNA sequences to create random sequences with the same nucleotide composition, yields similarly low structural complexities. Thus, the particular sequence composition of mitochondrial coding RNA sequences—i.e. low GC-content—is indeed sufficient to explain the observed simple secondary structures for coding RNA. Coding RNA is used in translation, and while secondary structure could still be relevant, e.g. to enhance stability (Mauger et al., 2019), it is likely not as critical as for noncoding RNA. Hence, coding RNA structures are well described by the sequence null model.

By contrast, noncoding RNA from the mitochondrion does not follow the same trend of low complexity. While situated in the same GC-poor genomic context, noncoding RNA genes feature higher GC-content (about 30%) than their surroundings, although this is still about 10% lower than nuclear RNA GC-content. These ncRNA maintain secondary structures with similar complexity as observed in the nuclear genome (see also Fig. S18).

Although at first sight the relatively high structural complexity might suggest positive selection, a more plausible explanation is that the mitochondrial ncRNAs originated in a different genomic context, similar to that of the nuclear genome, and have been preserved by purifying selection despite subsequent changes in their genomic context. The elevated GC-content of mitochondrial ncRNAs relative to the surrounding genomic environment may therefore reflect selective pressure to retain ancestral structural features. Alternatively, some mitochondrial tRNA genes may have been replaced by nuclear-encoded versions via horizontal gene transfer from other species (Salinas-Giegé et al., 2015).

### Simple and robust archaeal rRNA structures reflect environmental niches

Finally, we turn to ribosomal RNA (rRNA), the catalytic core component of the ribosome encoded by all life forms and therefore often featured in phylogenetic studies. We describe a similar analysis on tRNA in SI Section E. To investigate the distribution of rRNA phenotypes, we retrieved all reference sequences from SILVA (Quast et al., 2012), and used guide tree-based sampling for both the large subunit (LSU) and the small subunit (SSU) to obtain balanced samples with as much diversity as possible for the three domains and mitochondria and chloroplasts (see Methods). For SSU, sequence length varies considerably between the five subsets: archaeal SSU is smallest with many sequences shorter than *L <* 1000 nucleotides, whereas SSU from mitochondrial genomes is longest with some sequences of length *L >* 2000 (Fig. 5a). For LSU, sequence length distributions are more similar, with four subsets having means around *L* = 3000 and only eukaryotes featuring a higher mean as well as large variation around that mean (1500 ≲ *L* ≲ 4000; Fig. 5b). These observations suggest that the length of RNA molecules may be an important dimension for evolution.

**Figure 5.**
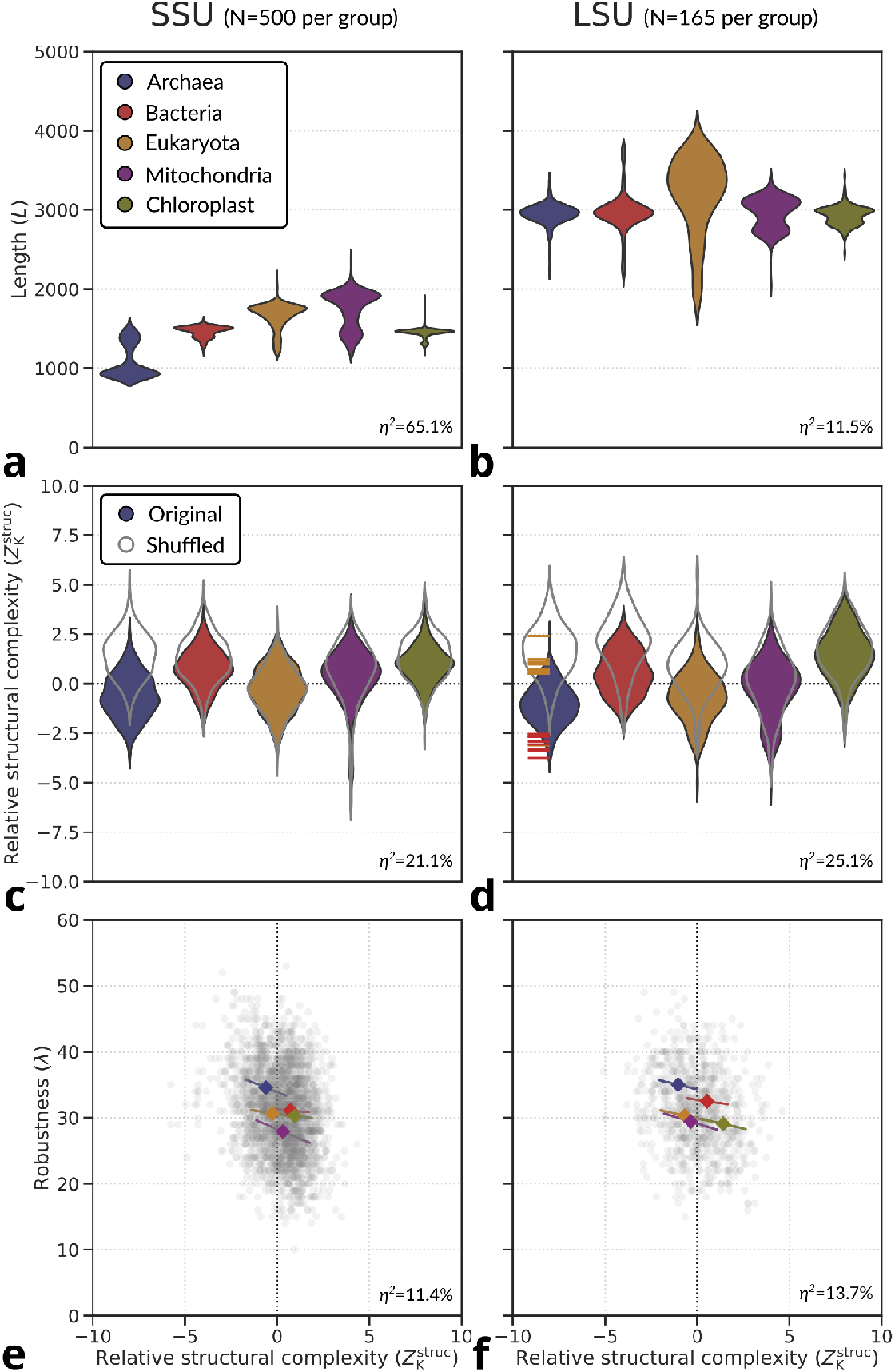
rRNA complexity distributions compared across taxa for (**a**–**b**) sequence length, (**c**–**d**) structural complexity and (**e**–**f**) robustness. ANOVA’s eta-squared statistic is shown at the bottom of each panel, indicating how much of the total variation is explained by the different taxa. In panels **c**–**d**, the structural complexity distributions for the shuffled version of the genes is shown with grey outlines. Red stripes in archaeal LSU rRNA denote annotated hyperthermophiles, and orange stripes thermophiles/mesophiles (see Table S3). (**e**– **f**) Robustness versus relative structural complexity. Individual rRNA molecules are shown in grey; diamonds display group means and line segments show a linear regression from one standard deviation below to one standard deviation above the mean on the x-axis. Standard errors are small and hidden behind the diamonds. Data was derived from the SILVA database (Quast et al., 2012). For the relation between mutational robustness and stability, see Fig. S19–21.

Despite substantial differences in sequence length, the relative structural complexities of the five subsets are similar, illustrating the utility of our complexity-versus-length scaling (Fig. 5c–d). Moreover, to first order they are all relatively close to the expected distribution from the sequence null model. To second order, there are small taxonomic differences. Compared to structures generated from random sequences, archaeal and eukaryotic rRNA molecules are structurally simple, whereas bacterial rRNA and rRNA from bacteria-derived organelles (mitochondria and chloroplast) are structurally more complex.

For many of the subsets, deviations from the sequence null model can be explained by GC bias, but interestingly, this does not work for archea. In paricular the structures of shuffled versions of archaeal rRNA, and to a lesser extent of bacterial rRNA, are more complex than structures generated from random sequences, suggesting that the sequences have non-random features beyond GC-content alone (Fig. 5c–d). Therefore, the relative simplicity of natural archaeal rRNA structures is all the more striking and could represent a signature of adaptation.

Simple phenotypes, which are typically encoded by many genotypes (Dingle et al., 2018; Johnston et al., 2022), tend to exhibit greater mutational robustness than complex ones. However, this correlation is not a physical necessity: some simple phenotypes are encoded by only a few genotypes (Dingle et al., 2020), and robustness can vary widely among genotypes mapping to the same phenotype (Aguirre et al., 2011; Mohanty et al., 2023). We evaluated this relationship computationally, defining as before the mutational robustness *λ* as the fraction of single-point mutants that retain the original secondary structure. Overall, we find only a weak correlation between robustness and relative structural complexity (Fig. 5e–f). For example, plastid rRNAs, particularly LSU, are structurally complex but more robust than mitochondrial or nuclear-encoded eukaryotic rRNAs. Among all groups, archaeal rRNAs are the most robust for both SSU and LSU, significantly exceeding the robustness of shuffled controls, which average *λ ≈*27.5.

Many archaea are extremophiles. Out of the archaeal species that are identified in our sample, those with very low structural complexity 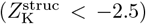 are almost exclusively hyperthermophiles, while those with relatively high structural complexity 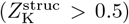 are acidophilic or mesophilic methane producers (Fig. 5; Table S3).

Although sequence length can confound stability comparisons, archaeal rRNAs are also the most temperature-robust among all subsets, in addition to being the most mutation-robust (Figs. S19–20). Together, these results suggest that archaeal rRNAs have been shaped, potentially by selection, toward a distinct set of phenotypes not typically sampled by random sequences. While the other subsets did not show as clear deviations from random, our analysis of archaeal rRNA suggests that with more fine-grained data, more subtle trends and small-scale adaptive signatures can potentially be elucidated. Nevertheless, compared to the full spectrum of physically possible RNA structures generated by the phenotype null model, most rRNAs deviate relatively little from the sequence null model.

## Discussion

To explore how evolution shapes molecular phenotypes, we analyzed the distribution of RNA secondary structures using a theoretically grounded complexity metric. Across a broad range of lengths and for nearly all taxonomic and functional categories, the structural distributions of noncoding RNAs closely match those generated by a simple null model that folds random sequences, accounting, where applicable, for non-random sequence features such as GC-content.

The close alignment between natural RNA and structures generated by random sequences, which simulates neutral evolution (Schaper and Louis, 2014; Johnston et al., 2022), is surprising. First, there is strong evidence that functional ncRNA is under purifying selection (Smith et al., 2013; Rivas, 2021; Walter Costa, 2023; Mattick et al., 2023; Backofen et al., 2024), implying a tight coupling between function and structure. Second, given RNA’s well-established evolvability (Schuster et al., 1994; Huynen, 1996; Schultes and Bartel, 2000; Greenbury et al., 2022; Srivastava et al., 2025) (SI, section F), we expected the diverse functional roles and environmental contexts in which ncRNAs evolved to have left more discernible imprints on their structural distributions.

Instead, the dominant global pattern reflects a strong developmental bias favoring simple structures. Notably, we find not merely a general tendency toward simplicity, but that folding a surprisingly small set of random sequences provides detailed predictions for the entire distribution of structural features seen in natural ncRNAs. For instance, for *L* = 300 ncRNA we find most of the diversity in about 10^5^ random samples, an unimaginably small fraction (Louis, 2016) of the 4^300^ ≈4× 10^190^ possible sequences. These results indicate that the structural repertoire of natural ncRNAs is highly constrained relative to the full space of potential folds.

These results also raise counterfactual questions about biological “dark matter”: Might the vast space of unexplored folds harbor novel functions? The success of artificial selection in uncovering numerous ribozymes absent from nature (Higgs and Muller, 2025), and function found in random sequences (Ekland et al., 1995; Neme et al., 2017), suggests that this hidden structural landscape could offer a rich source of functional innovation, particularly if search strategies can be directed beyond naturally favored regions. Our findings also speak to the longstanding debate between structuralism and functionalism in biology, which asks whether biological forms arise primarily from the demands of adaptive function or from underlying biophysical constraints (see Novick (2023) for a review). Here, we present an extreme case of the latter: a structuralist explanation shaped by inherent biases in the genotype-phenotype map.

Nevertheless, there is also little doubt that natural selection has shaped the emergence of functional ncRNAs (Ponting et al., 2009; Smith et al., 2013; Kapusta and Feschotte, 2014). So why are adaptive signatures in global secondary structure distributions so weak? One possibility is that secondary structure is not always central to function. This may hold, for example, for lncRNAs, where only portions of the structure tend to be conserved (Mattick et al., 2023). However, many functional RNAs do show clear evidence of strong purifying selection on structure (Smith et al., 2013; Rivas, 2021; Walter Costa, 2023; Backofen et al., 2024). For RNAs with defined structural roles, such as rRNAs, tertiary structure is critical, and while multiple tertiary folds may be consistent with a given secondary structure, the latter still imposes significant constraints (Vicens and Kieft, 2022), explaining its evolutionary conservation.

Another possibility is that RNA structure arose largely through non-adaptive processes (Lynch, 2007; Fernández and Lynch, 2011; Lukeš et al., 2011; Brunet et al., 2021; Torri et al., 2022) such as constructive neutral evolution (Stoltzfus, 1999). In this picture, functions can emerge spontaneously—for example, from the merger of two neutrally evolving structures—with selection acting only to preserve the new function. However, it would be surprising if such mechanisms dominated the evolution of functional RNA.

Alternatively, it could be that RNA is less evolvable than assumed, causing populations to become trapped at local optima near enough the original structures to not leave a clear structural imprint. However, this seems unlikely because the number of random sequences required to recover most of the structural diversity found in natural RNAs is orders of magnitude smaller than the number of distinct RNA molecules likely to have existed throughout natural history. If evolvability were limited in this way, we would expect to observe far greater variation in structural properties than is seen. While our sample sizes are too small to capture the full extent of functional diversity—which often requires additional sequence specificity (Higgs and Muller, 2025)—our results suggest that function may be easily accessible from a small set of structural scaffolds. We hypothesize that adaptation acts primarily on specific nucleotide subsets—such as ribozyme active sites—to generate function without requiring major structural changes. If correct, this suggestion that function is easy to find supports the RNA world hypothesis.

Our analysis has focused on first-order patterns in RNA structural distributions. We highlight one potential example of adaptation shaping global structure in extremophile archaea, though the signal is modest and may reflect indirect selection for robustness or thermal stability rather than direct optimization of structure. All this suggests that more detailed analysis and finer-grained structural metrics are needed to uncover additional signatures of adaptation, and to understand how it interacts with structure. In particular, if technical challenges can be addressed, extending this global analysis to RNA tertiary structures should uncover clearer evidence of adaptive shaping.

Finally, this work raises the question of whether similar global patterns exist in protein structures. Although adaptive changes in protein structure are well documented (Ortlund et al., 2007; Romero and Arnold, 2009; Szilágyi and Závodszky, 2000; Deng et al., 2010), neutral processes (Lynch, 2007; Fernández and Lynch, 2011; Lukeš et al., 2011; Brunet et al., 2021) and GP map biases (Herrera-Álvarez et al., 2025) also play important roles. It may also be valuable to examine other GP maps—particularly gene regulatory networks—which offer a promising framework for studying how evolution both shapes and is shaped by developmental bias.

## Methods

### Filtering of RNAcentral datasets

After downloading all RNA molecules of a specific length (*L* = 100 to *L* = 500) from RNAcentral, each dataset was subjected to two filtering steps (Fig. S10). First, only sequences which had no quality control warnings in RNAcentral were selected. There are four possible reasons for an RNA sequence in RNAcentral to be assigned a quality control warning: (1) potentially representing an incomplete sequence, (2) potential contamination (taxonomic mislabelling), (3) potential RNA type mislabelling, and (4) potentially representing coding RNA (i.e. containing an open reading frame). For *L* = 300, 2603 of 10383 (25.1%) bacterial sequences and 9117 of 14307 (63.7%) eukaryotic sequences passed this check. Second, we removed sequences that had more than 80% sequence identity with any other sequence in the dataset, such that the remaining data did not contain many copies of the same or closely related RNA molecules which would otherwise exacerbate database biases. For *L* = 300, only 292 bacterial sequences and 5159 eukaryotic sequences remained after this step, representing 2.8% and 36.1% of the original data, respectively. For both bacteria and eukaryotes, we observe that the complexity distributions shift towards slightly lower values after each filtering step (Fig. S10).

### Structural complexity of RNA using Lempel-Ziv compression

Structural complexity can be measured in various ways. Here, we adopt an approach from simplicity bias theory (Dingle et al., 2018), which uses compression-based complexity. The core idea is intuitive: structures with repeating patterns can be compressed and are considered simple. By contrast, structures lacking regular patterns appear random, are less compressible, and are considered complex. Thus, simplicity corresponds to compressibility, while complexity reflects incompressibility.

RNA secondary structures are routinely represented in dot-bracket form. We convert the dot-bracket string representing secondary structure to binary via the following: ‘.’ ‘→00’, ‘(‘‘→10’, ‘)’ ‘→01’. Thus, dot-bracket strings of length *L* are converted into binary strings of length 2*L*. Subsequently, we apply Lempel-Ziv compression (Lempel and Ziv, 1976) to the binary string *x* = *x*_1_…*x*_*n*_, calculating the number of dictionary elements *N*_*w*_(*x*_1_…*x*_*n*_) needed to store the string *x*, and averaging between forward and backward reading of the string (as in Dingle et al., 2018):

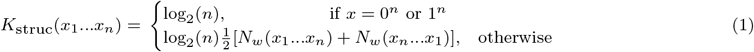

### Sampling from the phenotype null model

We use the method of Burghardt and Hartmann (2007) to effectively sample uniformly all secondary structures that are physically consistent with the rules of RNA folding. Although secondary structures sampled this way may not actually be designable representing the minimum-free-energy state of any sequence, (Dingle et al., 2015) found that the distribution of the numbers of stacks is very similar when derived from computational random sampling, and when using analytical mathematical methods (see also Ghaddar and Dingle, 2023). This alignment indicates that the mathematical results and computational sampling are likely to be consistent.

### Diversity sampling of rRNA guide trees from SILVA

To allow for fair comparisons, we down-sampled the enormous amount of data in SILVA to equal numbers for each of our five subsets: archaea, bacteria, eukaryotes, mitochondria and chloroplasts. These five subsets correspond to five root nodes in the SILVA guide trees, which we split into five separate trees. Then, we removed leaves constituting metagenomic samples and leaves whose taxonomic classification was in one of the other subsets (e.g. bacterial rRNA in the mitchondria subtree). Furthermore, we had to remove sequences that included faulty characters (e.g. undefined nucleotides) and we threw out taxa for which we were unable to find genomic information in NCBI (genome size and GC-content). At the end, the mitochondrial subtree contained the fewest leaves for both SSU (*N* = 587, rounded down to *N* = 500) and LSU (*N* = 165). For each of the other subtrees, we obtained the number of branches present when cutting the tree at 0.01 distance intervals starting from the root. We cut a tree at the first instance that the number of branches is more than the desired value (*N* = 500 for SSU, and *N* = 165 for LSU). Since the number of branches is usually not exactly the target value, we randomly remove branches until we hit our target. From each remaining branch in the pruned tree we then randomly pick one of the terminal leaves, i.e. a single rRNA gene. In sum, this method ensures that we sample from each subset as much phylogenetic diversity in rRNA genes as possible.

## Acknowledgements

We thank Zachary Ardern and Paulien Hogeweg for helpful conversations, and Bernard Kuhle for providing filtered tRNA data. NSM acknowledges support of the Spanish Ministry of Science and Innovation through the Centro de Excelencia Severo Ochoa (CEX2020-001049-S, MCIN/AEI/10.13039/501100011033), the EMBL partnership and the Generalitat de Catalunya through the CERCA programme. This research is part of grant JDC2022-049526-I to NSM, funded by MCIU/AEI/10.13039/501100011033 and by “European Union NextGenerationEU/PRTR”.

## Code Availability

Our user-friendly tool for calculating structural complexity and sequence complexity of any RNA molecule is available online: https://colab.research.google.com/drive/1VaT3N Ga Tr0Bt47oYfldYy oaJnWR5 v?usp=sharing

