## Supplementary Materials for "RNA secondary structures are conserved but random"

August 18, 2025

#### Supporting Information Text

##### A. The complexity distribution of RNA structures upon uniform sampling of sequences

In the main text, we follow Dingle et al. Dingle et al. (2018) and use a Lempel-Ziv compression algorithm to quantify the structural complexity of RNA secondary structures, represented in dot-bracket notation. Simple structures, such as `...................` or `..........(...)..........`, can be highly compressed and thus yield low values of $K_{\text{struc}}$. In contrast, more complex structures, such as `....((..))..((((...))))..(..)..`, are less compressible and exhibit higher $K_{\text{struc}}$ values.

This definition prompts a natural question: what is the expected distribution of structural complexities if RNA sequences are sampled uniformly at random (that is the probability of picking any sequence is independent of the sequence) and folded into their minimum free-energy secondary structures?

###### A.1 Simplicity bias for individual structures

On the one hand, previous work Dingle et al. (2018); Johnston et al. (2022) has shown that the RNA genotype–phenotype (GP) map exhibits a phenomenon known as simplicity bias. What this means is that when sequences are sampled uniformly at random, simple structures (with low complexity) tend to occur much more frequently, while individual complex structures appear much less often. That is, the RNA GP map is biased toward simpler phenotypes.

This bias has broad theoretical foundations in algorithmic information theory Dingle et al. (2018, 2020). It can be intuitively understood in terms of how genetic mutations explore a space of developmental programs (algorithms) encoded by the GP map Johnston et al. (2022); Martin et al. (2024): A random search in the space of algorithms is more likely to find shorter algorithms than longer ones, and structures that can be described by shorter algorithms are simpler.

A central prediction of this framework is that the probability $Pr(P)$ of observing a phenotype $P$ is upper-bounded by $2^{-K(P)}$, where $K(P)$ is the Kolmogorov complexity of $P$. Empirical studies show that this upper bound is often well approximated by $Pr(P) \approx 2^{-a\tilde{K}(P)+b}$, where $a$ and $b$ are constants, and $\tilde{K}(P)$ is a compression-based estimate of $K(P)$. In our context, $\tilde{K}(P)$ corresponds to $K_{\text{struc}}(P)$. Simplicity bias is illustrated schematically in Fig. S1a, where the upper bound defines an exponential drop-off in $Pr(P)$ as complexity increases linearly, with most of the total probability mass near this bound.

###### A.2 Number of structures versus complexity

On the other hand, although simple structures are more probable individually, they are also fewer in number. By contrast, complex structures are more numerous, though individually less probable. This relationship can be quantified using binary strings, where it can be easily shown that the number of strings with Kolmogorov complexity $K$ scales as $2^K$, illustrated by the brown line in Fig. S1b.

If the simplicity bias is exact (i.e., $Pr(P) = 2^{-K(P)}$), these two effects perfectly balance, resulting in a uniform distribution $Pr(K)$ over complexities, as shown in Fig. S1c. A similar scaling behavior has been observed in deep neural networks, where the probability of producing a function $f$ of complexity $K(f)$ scales as $2^{-K(f)}$ when network weights are sampled at random Mingard et al. (2025). The analogy to the GP map arises when one thinks of the weights as the genotypes, and the functions as the phenotypes.

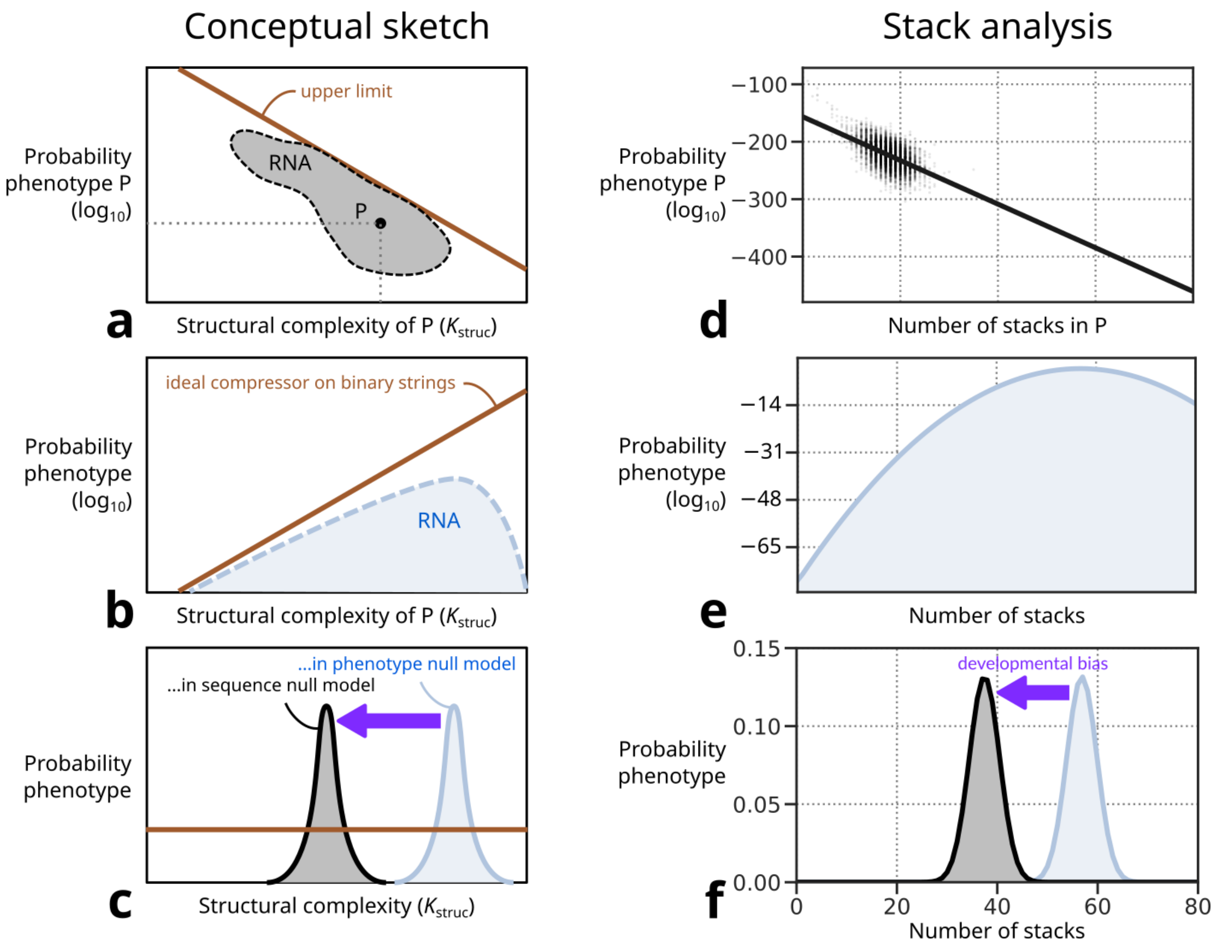


Figure S1 – Schematic of the complexity distributions and how they interact. (**a**) A sketch of the probability of obtaining an individual phenotype P as a function of complexity $K_{\text{struc}}$ upon random sampling of sequences shows simplicity bias. This figure is modeled on Fig. 1a of Dingle et al. (2015) which shows this distribution for RNA molecules of length $L = 55$. Simplicity bias means that high probability phenotypes have low complexity, and phenotypes with higher complexity have exponentially fewer genotypes mapping to them. Most of the probability weight is near the upper-bound given by $2^{-aK_{\text{struc}}+b}$. (**b**) For an ideal compressor that can perfectly compress strings, the number of binary strings with complexity $K$ scales as $2^K$. For RNA structures, the number of phenotypes with complexity $K_{\text{struc}}$ should scale exponentially with complexity for most of the complexity range, but will deviate for low and high complexities. (**c**) The complexity distribution when sampling random genotypes (black) has a substantially lower mean than the complexity distribution when sampling random phenotypes (light blue), showing the effect of phenotypic bias (note that the y-axis is much smaller in (c) than in (b). By contrast, for an ideal compressor on binary strings, the complexity distribution will be flat. (**d**) Correlation between the frequency of a phenotype and its number of stacks for natural RNA molecules (dots, $N = 6450$) and structures produced by sampling random sequences of length $L = 300$ (black line). The relative frequency in sequence space was estimated using the code and method described in Weiß and Ahnert (2020). A linear model was then fitted to this log-linear plot (Pearson's correlation coefficient $r = 0.54$). (**e**) Distribution of number of stacks among random phenotypes as extrapolated from the analytical normal distributions for $L = [10, 30]$ (Hofacker et al., 1998) to $L = 300$. (**f**) The sequence null model and phenotype null-models for stacks are shown in grey and blue, respectively. The mean and standard deviations of these analytic normal distributions were fit with high precision ($R^2 = 99.99997\%$ and $R^2 = 99.98\%$, respectively) although being limited to a very small range due to rapidly increasing computational costs with sequence length.

The reason there are many more possible secondary tructures with high complexity than with low complexity is combinatorial, in that there are few ways to be simple (e.g., `..........`) but many ways to be complex (e.g., `((...)..).` or `..((..)..))`). Since dot-bracket notations can be mapped to binary or ternary strings, one might expect the number of structures to scale exponentially with complexity. However, constraints inherent to RNA folding—such as the requirement that every open bracket be closed—mean that highly complex or extremely simple strings may not correspond to physically valid RNA structures.

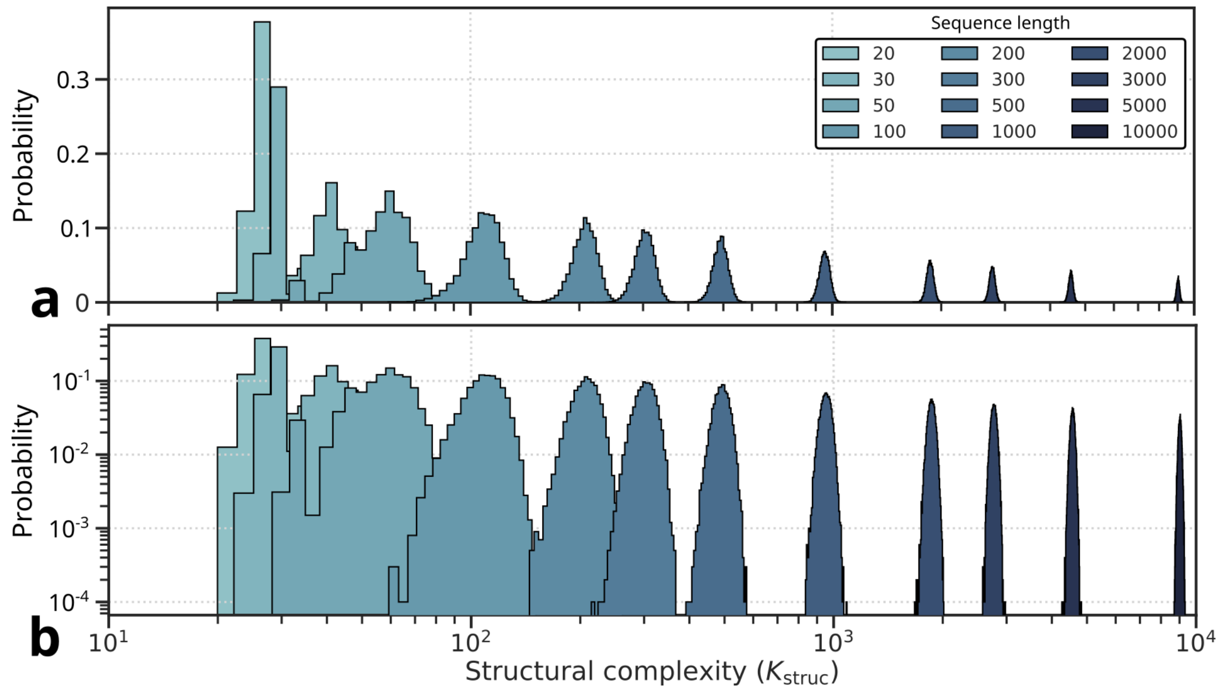


Figure S2 – Probability distributions of structural complexity when sampling random genotypes and folding them, for lengths $L = [20, 10000]$ on a (**a**) linear scale and on a (**b**) logarithmic scale. At each length, a sample of fixed size ($N = 10^4$) was taken. Histograms show one bin per complexity level (complexity is quasi-discrete and the step size scales with sequence length).

At the higher complexity end, this argument helps explain why Fig. S9 shows that the distribution of complexities for all valid phenotypes peaks at a lower value than the distribution for all possible ternary strings.

Similarly, simple strings such as ((((((((((((((( or ))))))))))))))) are invalid due to unmatched parentheses. Thus, both ends of the complexity spectrum are constrained: very complex and very simple structures are restricted by folding rules and physical feasibility. As a result, the actual distribution deviates from the idealized $2^K$ growth, as suggested by the putative blue curve in Fig. S1b.

##### A.3 Combining the two effects for the RNA GP map

When sampling random phenotypes, only the upper portion of the putative distribution is accessible (note that Fig. S1b is plotted on a logarithmic scale), resulting in a narrow distribution of high-complexity values (illustrated in Fig. S1c). Empirically, we find that these sampled phenotypes follow an approximately normal distribution in $K_{\mathrm{struc}}$. Further analysis of this distribution presents an interesting direction for future work, but is not needed for our current investigation.

By contrast, when we sample random sequences and fold them—our sequence null model—the observed distribution of structural complexity, $Pr(K)$, reflects the product of two components: the distribution of complexity across all phenotypes (light blue curves in Fig. S1b), and the frequency with which individual phenotypes arise from random genotypes (Fig. S1a). At the scale of our sampling, this results in a structural complexity distribution that is also approximately normal (schematically in Fig. S1c and for lengths $20 \leq L \leq 10000$ in Fig. S2), but with a mean shifted to significantly lower complexity values compared to the random phenotype distribution. We show the difference between the random phenotype distribution and that produced by the sequence null model in Fig. 1 in main text for $L = 300$, and for lengths $30 \leq L \leq 500$ in Fig. 3 of the main text, revealing that the differences in the means grow with length. In other words, due to simplicity bias, the sequence null model recovers a highly biased subset of relatively low—but not the lowest—complexity phenotypes.

A good simpler proxy for complexity $K_{\mathrm{struc}}$ is the number of stacks Dingle et al. (2015, 2018); Johnston et al. (2022). We uncover a similar trend in the distributions of the number of stacks as seen for the different $K_{\mathrm{struc}}$ distributions (Fig. S1d–f). The advantage here is that there are analytic approximations that can be used for some of the steps, providing further support for our results.

#### B. Complexity distributions for sequences

Compression-based techniques have long been used to analyze biological sequences, including RNA Grumbach and Tahi (1994); Li et al. (2004); Giancarlo et al. (2009). While we don't pursue this direction much in this paper, it is useful to briefly summarize how our Lempel-Ziv based sequence complexity measure $K_{\mathrm{seq}}$ reflects the compressibility of a sequence. For example, highly repetitive sequences yield lower $K_{\mathrm{seq}}$ values, while random, patternless sequences are less compressible and thus have higher values. Note that this measure goes beyond entropy. For example, $ACGUACGUACGUACGUACGUACGUACGU$ is a maximum entropy sequence, but it would have a low $K_{\mathrm{seq}}$ due to its repetitive nature. As shown in the supplementary materials of Dingle et al. Dingle et al. (2018), LZ complexity follows a $2^K$ distribution for low $K$, and approximates a Gaussian for higher values, which encompasses most RNA sequences. While it is imperfect, we will use a Gaussian approximation to the sequence complexity to quantify our measure of how far a sequence deviates from the expected mean $K_{\mathrm{seq}}$.

#### C. Verifying normality of complexity distributions

We observed in the plots above that the complexity distributions we measured are approximately Gaussian. Instead of relying on statistical tests, we assess the distributions with Q–Q plots, comparing the quantiles of data points in our measured distribution with the quantiles that they would have if drawn from a normal distribution with the same mean and standard deviation. Fig. S3 shows that the samples resemble well-behaved normal distributions within roughly four standard deviations below and above the mean. The remaining deviations from normality are largely due to the discreteness of the complexity values. Note that the highest probability function is always the simplest one with no bonds. We exclude this structure from our Gaussian fits. It is important to note that we cannot rule out that the distributions are non-Gaussian upon further sampling; this has to be explicitly measured, but it is computationally expensive. We use our Gaussian approximation as a rough guide to extend the distribution. Even if this extension is not perfect, which we expect to be the case, it can still be usefully employed as a measure of how far one deviates from the mean value.

#### D. Reliability of RNA structure predictions

We used the ViennaRNA package to calculate the minimum-free-energy structure for each natural and random sequence. Such free-energy-based methods are thought to be accurate—a closely related method correctly predicts $73 \pm 9\%$ of canonical base pairs for a database of known RNA structures up to lengths of 700 nucleotides (Mathews et al., 2004)—and are expected to work best for shorter RNA (Reuter and Mathews, 2010), such as analyzed in Dingle et al. (2015, 2022). Dingle et al. (2022) indeed showed that coarse-grained RNA shapes are similar between random and natural RNA of length $L \approx 100$ independently of whether their structural data was based on ViennaRNA predictions or on alignment to the consensus structure of the seed alignment for respective RNA families. Furthermore, Ghaddar and Dingle (2023) showed that structural motif trends in RNA secondary structure as predicted by ViennaRNA are consistent across length scales from $L \approx 100$ to $L \approx 3000$. Thus, free-energy-based structure prediction methods are suitable for this kind of analysis. In addition, we performed a direct validation of our findings by analyzing experimentally refined secondary structures as extracted from the Rfam database (Fig. S4; ref Ontiveros-Palacios et al., 2024). Like most predicted RNA secondary structures, the bulk of experimental data has a relative structural complexity $Z_K^{\mathrm{struc}}$ close to 0, resembling random genotypes and supporting our conclusions based on ViennaRNA predictions.

#### E. Absence of domain-level trends in complexity in tRNA genes

Similar to the analysis of two rRNA genes, we also studied structural complexity for 20 transfer RNA types (Fig. S5). tRNA and rRNA not only fulfill different cellular functions, but they also represent the two extremes of the variation in sequence length: most tRNA molecules are about 70 nucleotides long, whereas rRNA molecules typically span several thousand nucleotides. Perhaps because they are so small, there is limited variation in length in tRNAs between domains, except for mitochondrial tRNA which in many cases is shorter. Previous work has shown that tRNAs in the mitochondria of bilaterians are adapted to the higher mutation rates and shortening of sequences through the evolution of a different recognition mechanism between tRNA and cognate aminoacyl–tRNA synthetase (Kuhle et al., 2020). Nevertheless, we do not observe substantial differences in relative structural complexity between mitochondrial tRNA and nuclear eukaryotic tRNA or tRNA from bacteria or archaea, for any of the 20

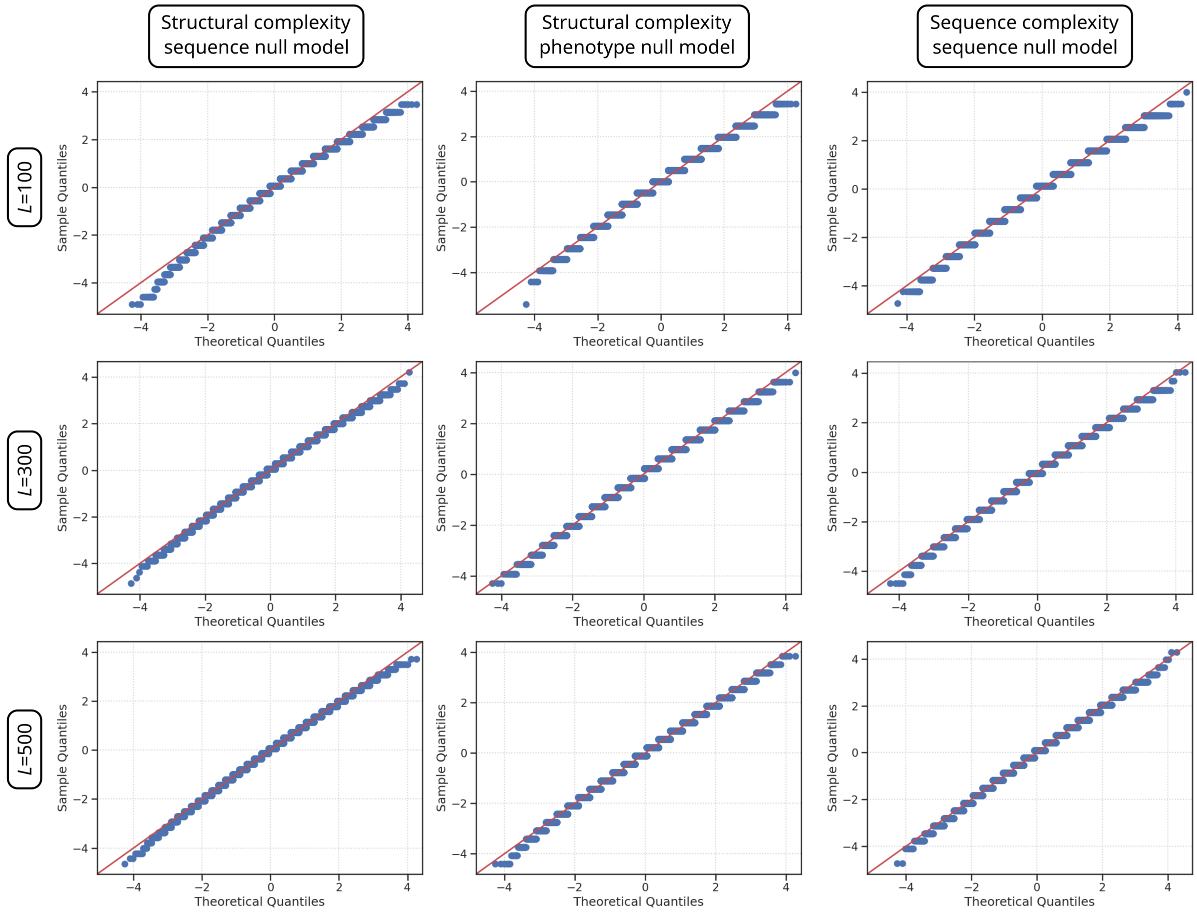


Figure S3 – Q–Q plots for structural complexity ($K_{\text{struc}}$) of structures generated by our sequence null model and our phenotype null model. We also show sequence complexity ($K_{\text{seq}}$) of random genotypes at three different lengths showing minimal deviations from normality (i.e. distance to the red diagonal). Note that the full distribution of LZ-based sequence complexity will deviate from Gaussianity, especially for lower complexities (see Supplementary Figure 2 in Dingle et al. Dingle et al. (2018)). Nevertheless, we can use these Gaussian fits to measure how far a sequence deviates from a fully random sequence.

types (the shortness of mitochondrial tRNA seems to be the only sense in which it is simpler than other tRNA).

In general, there is considerable variation in relative structural complexity within each subset but there are no clear differences between subsets. Even in mitochondrial genomes, we find both tRNA with high and tRNA with low relative structural complexity. We hypothesize that genome-wide trends such as different biases in GC-content between different datasets do not affect tRNA genes as much, because unlike the much longer rRNA genes, selection can more easily oppose these biases in short tRNA sequences. On the other hand, the large variation within each dataset suggests that there is a substantial degree of functional neutrality among tRNA phenotypes. Out of all variation in relative structural complexity, tRNA type explains 10.0% and genus explains 9.3%. It would be interesting to investigate these differences further to see if, for example, a signature of selection could be found at the genus level.

#### F. Simulated directed evolution of RNA towards rare structures

The absence of signatures of positive selection in natural RNA structures could indicate that selection for structure is too weak to overcome the enormous developmental bias. However this hypothesis contradicts previous theoretical work which has showcased that RNA is very evolvable (Schuster et al., 1994; Huynen, 1996; Schultes and Bartel, 2000; Greenbury et al., 2022; Srivastava et al., 2025). To test these prior results, we perform evolutionary simulations where we simulate discrete generations of a population of 100 RNA molecules of length $L = 100$. For each new generation, molecules are selected proportional to their

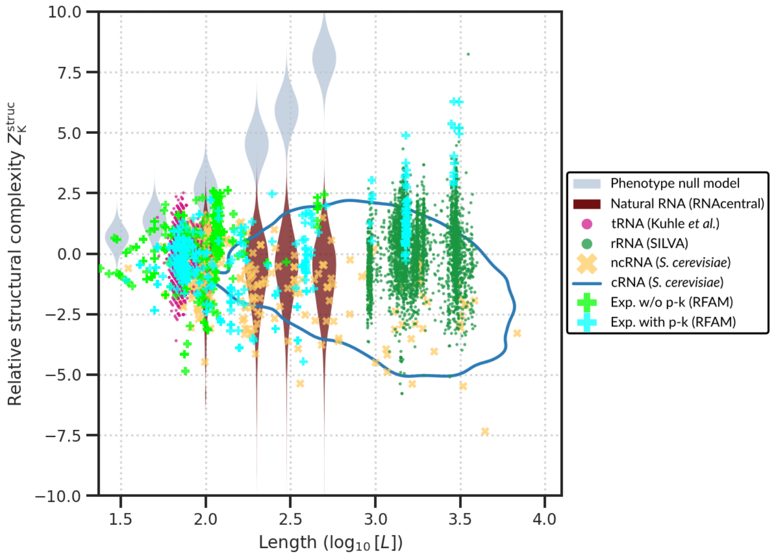


Figure S4 – Experimentally refined secondary structures overlaid on those predicted by ViennaRNA (Fig. 3). Three-dimensional experimental structures were obtained from Rfam (Ontiveros-Palacios et al., 2024) and secondary structures extracted using DSSR (Lu et al., 2015) and split into a set that contained no pseudo-knots (w/o p-k) and one that did contain pseudo-knots (with p-k). The pseudo-knots were removed from the latter set. In addition, structures without base pairs were ignored because the scaling relationship was not fitted to them.

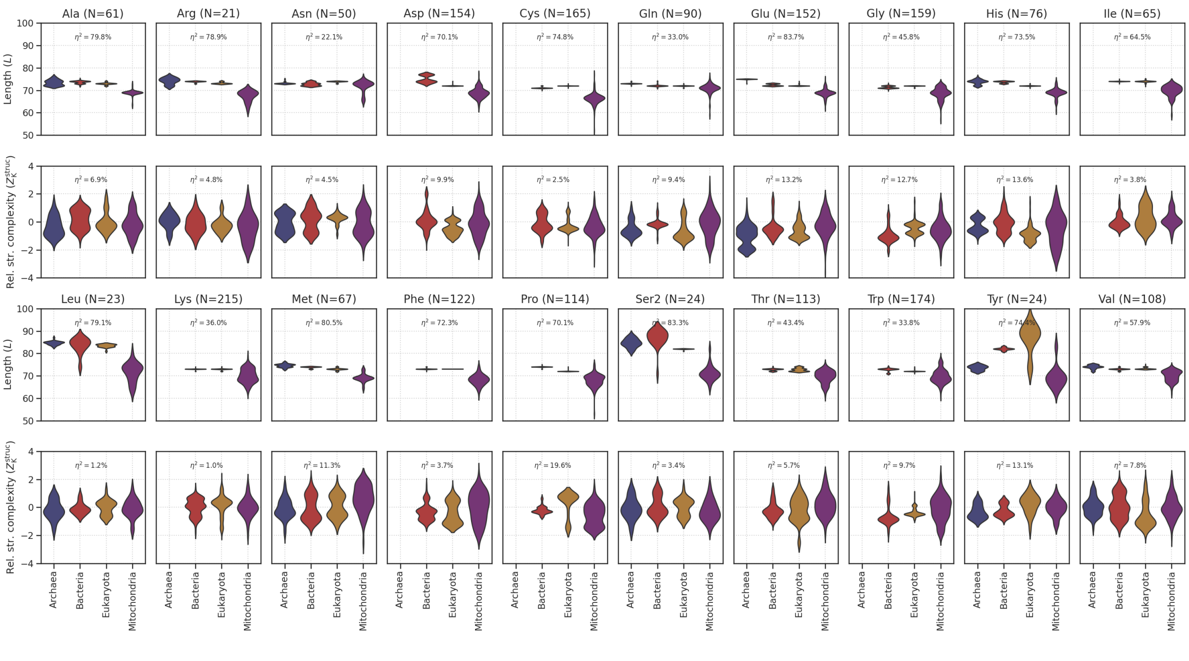


Figure S5 – Comparison of tRNA complexities between different domains and nuclear and mitochondrial genomes of eukaryotes, based on the data of Kuhle et al. (2020). Archaeal data was not available for all tRNA types. In all cases, datasets were downsampled to the same size as the smallest dataset, which is usually Archaea (if present), and otherwise Bacteria.

fitness (selection coefficient $s = 0.1$, i.e. $f = \varepsilon^{0.1K}$) and with mutations (substitutions) occurring at a rate of $\mu = 0.01$ per nucleotide (yielding on average one mutation per molecule per generation). We also explored other parameter values for population size, selection coefficient and mutation rate, but their quantitative effect does not change our main conclusion, namely that positive selection can easily find rare secondary structures in the full morphospace and that this leaves a clear signature on the structural complexity distribution of natural RNA.

We first run three sets of evolutionary simulations: (1) without selection, (2) selecting for low struc-

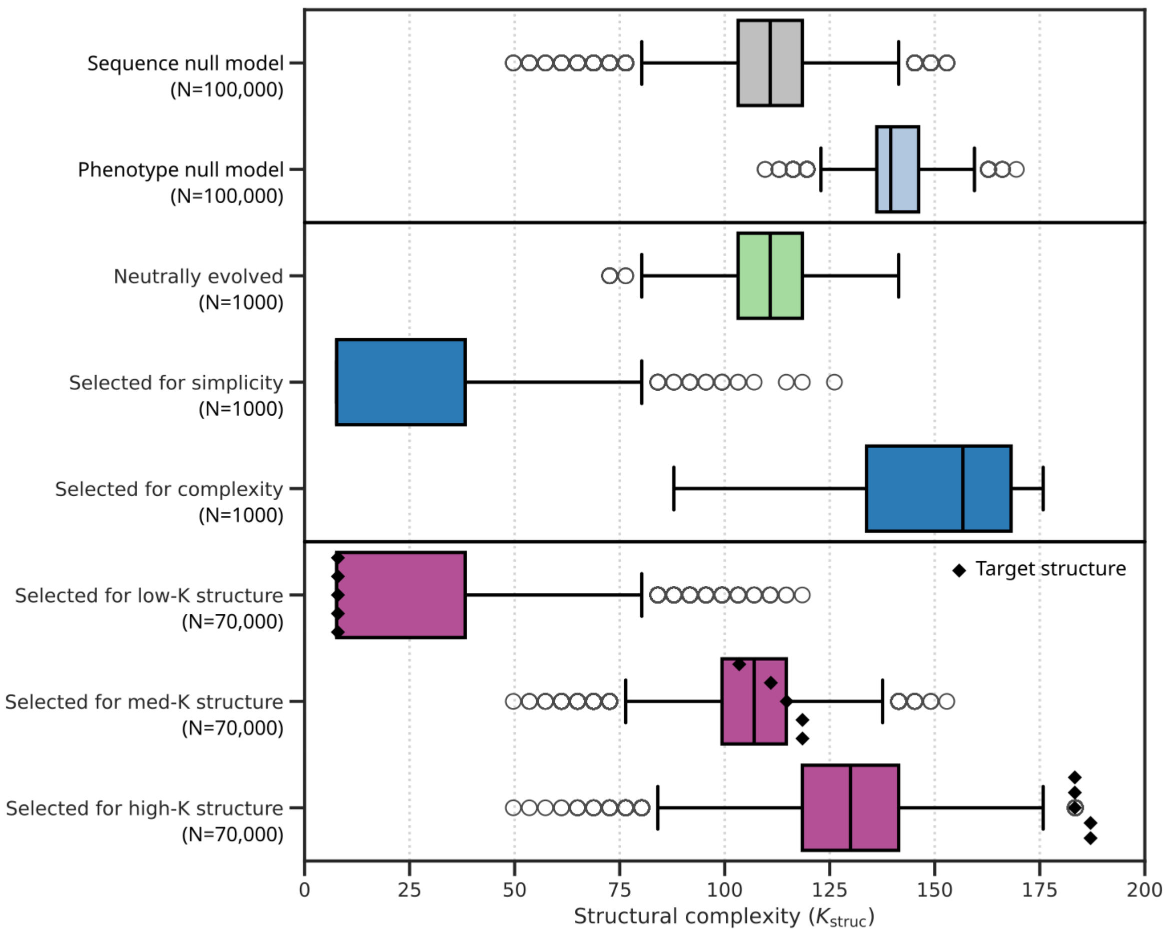


Figure S6 – Structural complexity distributions of RNA molecules after directed evolution compared to random genotypes and random phenotypes. Directed evolution yields rare structures with extreme structural complexity compared to random genotypes which have not evolved. Simulation parameters: $N = 100$, $s = 0.1$, $\mu = 0.01$, $t = 1000$. For the top three simulated datasets (green and blue), 10 runs are shown per experiment (5 initial molecules times 2 replicates). For the bottom three simulated datasets (purple), 700 runs are shown per experiment (14 initial molecules times 5 targets times 10 replicates).

tural complexity, and (3) selecting for high structural complexity. Neutrally evolving populations of RNA molecules (i.e. without selection) end up with a distribution of structural complexity that matches that of the sequence null model (Fig. S6). Earlier work already indicated that the phenotype distribution of randomly sampled sequences matches the result of neutral evolution Schaper and Louis (2014); Johnston et al. (2022), even when small insertions and deletions are taken into account Martin and Ahnert (2021).

In this work, we focused on the minimum-free-energy (MFE) fold as is customary in analyses of RNA folding. In reality, RNA folding is dynamic, represented by an ensemble of low-energy conformations (Vicens and Kieft, 2022). Still, recent work Dingle et al. (2022) indicates that taking suboptimal structures into account does not meaningfully change the phenotype distribution of randomly sampled sequences.

Unlike neutrally evolving populations, those under selection for either low or high structural complexity readily attain extreme values well beyond the range observed upon folding randomly sampled genotypes (Fig. S6). This result aligns with prior work demonstrating the high evolvability of RNA Schuster et al. (1994); Huynen (1996); Schultes and Bartel (2000); Greenbury et al. (2022); Srivastava et al. (2025). Under direct selection for complexity, developmental bias can be readily overcome, allowing access to rare phenotypes and producing structural complexity distributions that diverge markedly from those of the sequence-based null model.

Direct selection on structural complexity lacks a clear biological rationale. More realistically, under the sequence–structure–function paradigm, specific secondary structures may be selected for their functional roles. To model this, we conducted evolutionary simulations in which RNA fitness was defined by structural similarity to a predefined target. Similarity was quantified using the global Needleman–Wunsch alignment score between dot-bracket notations, with match = 1, mismatch = 0, and gap = −1 (Needleman and Wunsch, 1970).

This setup generates a smooth phenotype–fitness landscape, consistent with prior models, although recent studies suggest that even random landscapes can be traversed effectively by RNA (Greenbury et al., 2022; Srivastava et al., 2025). As targets, we selected 15 RNA molecules from prior simulations: 5 with median complexity (med-K), 5 with the lowest complexity (low-K), and 5 with the highest (high-K). Each of these was used both as a target and as a starting point for evolution, yielding 15 × 14 unique target–start pairs, each replicated 10 times.

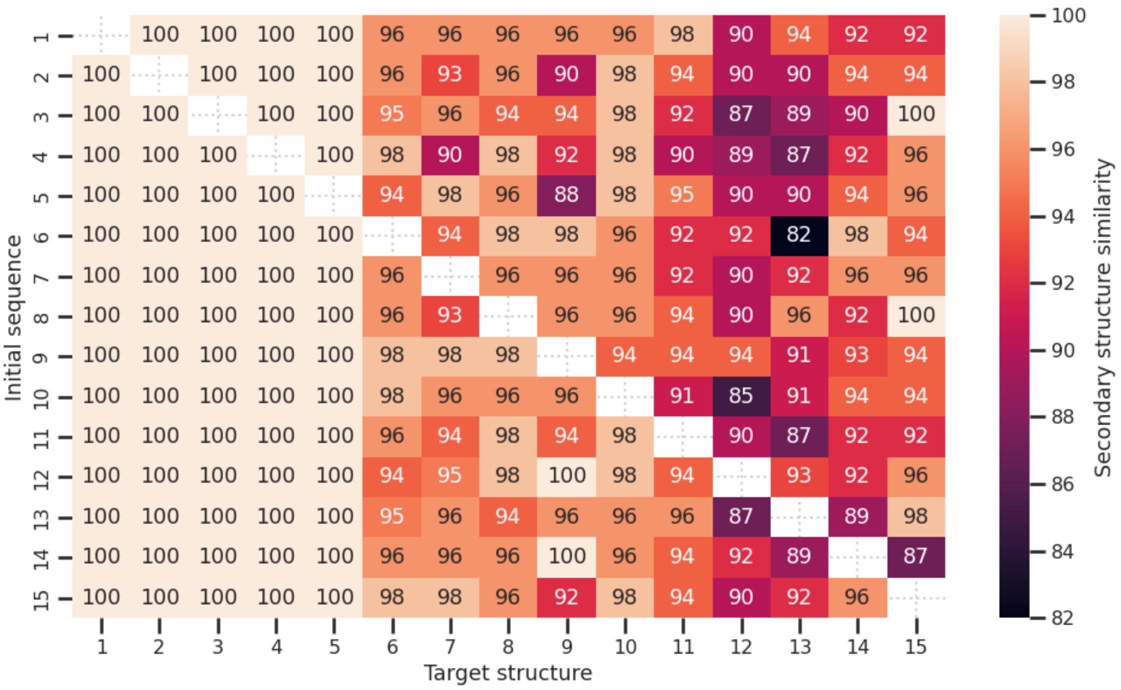


Figure S7 – Evolutionary success of populations evolving from a specific sequence towards a specific target structure. Each colored square shows the individual with the highest fitness (i.e. highest structural similarity to target) of 10 replicate populations of size $N = 100$ after $t = 1000$ generations. RNA sequences 1–5 all have the simplest possible structure (low-K); molecules 6–10 are random genotypes (med-K); molecules 11–15 have high structural complexity (high-K, see Fig. S6).

Even within just 1,000 generations, many populations evolve structures close to the target, with some reaching it exactly (Fig. S7). Evolutionary success is largely independent of the initial sequence but strongly influenced by the structural complexity of the target. The simplest target—the unfolded structure with no base pairs—is frequently discovered, and the resulting structural complexity distribution closely matches that of populations under direct selection for simplicity (Fig. S6). This is consistent with previous work showing that the unpaired structure is the most frequent one, and widely distributed in sequence space Dingle et al. (2015), making it readily accessible to evolutionary search.

By contrast, selecting for a high-complexity structure that is rare in sequence space is more difficult; most simulations failed to recover the target structure within 1,000 generations. However, high-fitness structures that are structurally similar to the target tend to exhibit comparable levels of complexity, producing a clear signature of selection in the structural complexity distribution. This suggests that even without direct selection on complexity—or even in the absence of the exact optimal phenotype—selection can leave a detectable imprint due to the structural similarity of evolved phenotypes to the target. These results reinforce earlier findings that RNA is highly evolvable, even in the presence of strong phenotypic bias.

### Supporting Figures and Tables

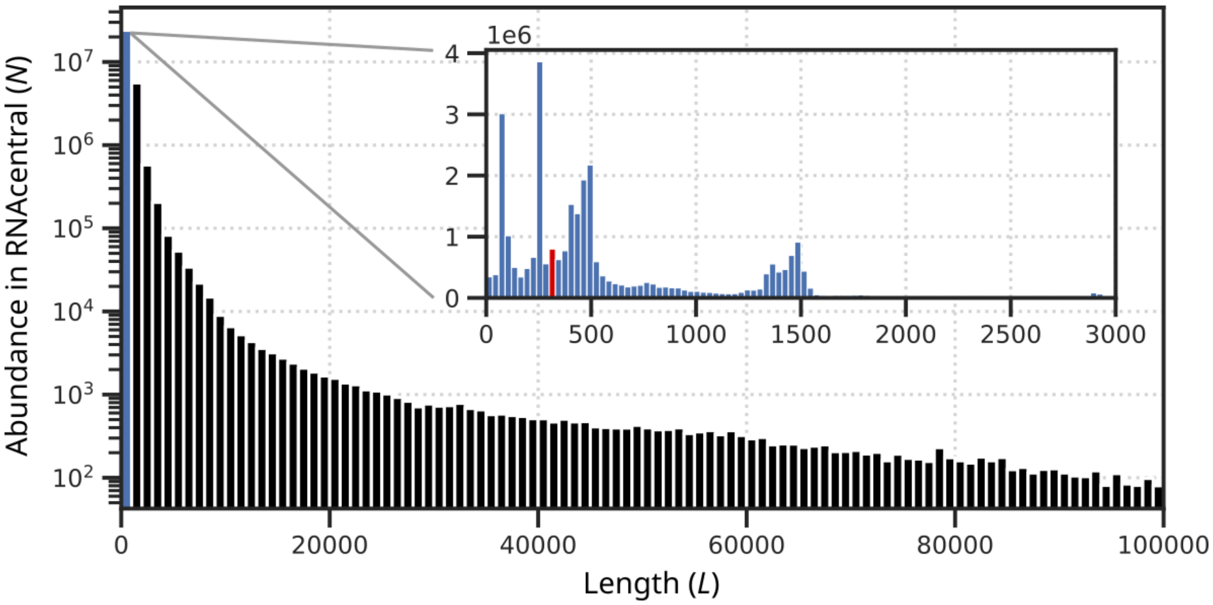


Figure S8 – Length distribution of sequences in RNAcentral after removing double entries ($N = 30,721,364$). The dataset covers lengths from $L = 10$ to $L = 583,414$. The main panel shows a large part of the variation on a logarithmic scale, with the inset showing abundances on a linear scale for lengths between $L = 0$ and $L = 3000$. The red bar highlights the interval that contains $L = 300$. Note that the data shown has not been quality controlled, or filtered by sequence similarity. As a result, sharp peaks (e.g. for $L \approx 250$) may not properly reflect the natural length distribution. The first sharp peak (around $L = 70$) represents abundant tRNA in the database, and the humps at $L \approx 1500$ and $L \approx 3000$ represent SSU rRNA and LSU rRNA, respectively (see Fig. 5 in main text).

Table S1 – Statistics for RNA molecules with extreme complexity values (sorted by structural complexity; see Fig. S11; turqoise markers in Fig. 1c in the main text).

| Label | RNA type | Species (common name) | Domain | $K_{seq}$ | $H$ | GC-content | $K_{struc}$ |
|---|---|---|---|---|---|---|---|
| URS000280CEAF | misc_RNA | *Melitaea cinxia* (Glanville fritillary) | Eukaryota | 161.5 | 1.55 | 0.33 | 9.2 |
| URS0001C7C540 | lncRNA | *Paramuricea clavata* (Violescent sea-whip) | Eukaryota | 230.7 | 1.51 | 0.49 | 46.1 |
| URS0002424CDB | lncRNA | *Brassica napus* (Rapeseed) | Eukaryota | 230.7 | 1.61 | 0.35 | 64.6 |
| URS0002388837 | lncRNA | *Elaeis guineensis* (African oil palm) | Eukaryota | 521.4 | 1.64 | 0.47 | 78.4 |
| URS000195249D | lncRNA | *Monodelphis domestica* (Gray short-tailed opossum) | Eukaryota | 286.1 | 1.37 | 0.38 | 83.1 |
| URS000234F001 | lncRNA | *Elaeis guineensis* (African oil palm) | Eukaryota | 558.3 | 1.65 | 0.49 | 83.1 |
| URS00026EB4CE | rRNA | *Serratia marcescens* | Bacteria | 632.2 | 1.99 | 0.56 | 83.1 |
| URS000241975B | lncRNA | *Physcomitrella patens* (Spreading earthmoss) | Eukaryota | 433.8 | 1.66 | 0.26 | 96.9 |
| URS0000C2B725 | sRNA | *Eimeria necatrix* | Eukaryota | 147.7 | 1.94 | 0.63 | 193.8 |
| URS0000CD0EA8 | rRNA | *Prevotella spp.* | Bacteria | 641.4 | 1.87 | 0.60 | 364.5 |
| URS0000CFCED9 | rRNA | *Alicyclobacillus spp.* | Bacteria | 632.2 | 1.86 | 0.58 | 364.5 |
| URS0000D478F8 | rRNA | *Yersinia spp.* | Bacteria | 655.2 | 1.92 | 0.57 | 364.5 |
| URS0000D4D582 | rRNA | *Branchiibius spp.* | Bacteria | 632.2 | 1.92 | 0.55 | 364.5 |
| URS0000E72498 | lncRNA | *Junco hyemalis* (Dark-eyed junco) | Eukaryota | 632.2 | 2.00 | 0.54 | 364.5 |
| URS0001A21C2E | lncRNA | *Oncorhynchus tshawytscha* (Chinook salmon) | Eukaryota | 332.2 | 1.90 | 0.44 | 373.8 |

Table S2 – Using ANOVA to detect differences in structural complexity between various subsets of RNA of length $L = 300$. There are 7 RNA types with $N > 50$: lncRNA, miscRNA, rRNA, snRNA, sRNA, SRP-RNA, RNase-P-RNA. Coeff. of var.: coefficient of variation; Corr. for seq. comp.: corrected for sequence composition.

| Grouping | Sample size ($N$) | Coeff. of var. ($\eta^2$) | Corr. for seq. comp. ($\eta^2$) |
|---|---|---|---|
| Natural vs random | 684 (228 per group) | 3.0% | 0.042% |
| Bacteria vs eukaryota | 456 (228 per group) | 11.9% | 0.074% |
| RNA types (x7) | 357 (51 per group) | 14.2% | 4.8% |

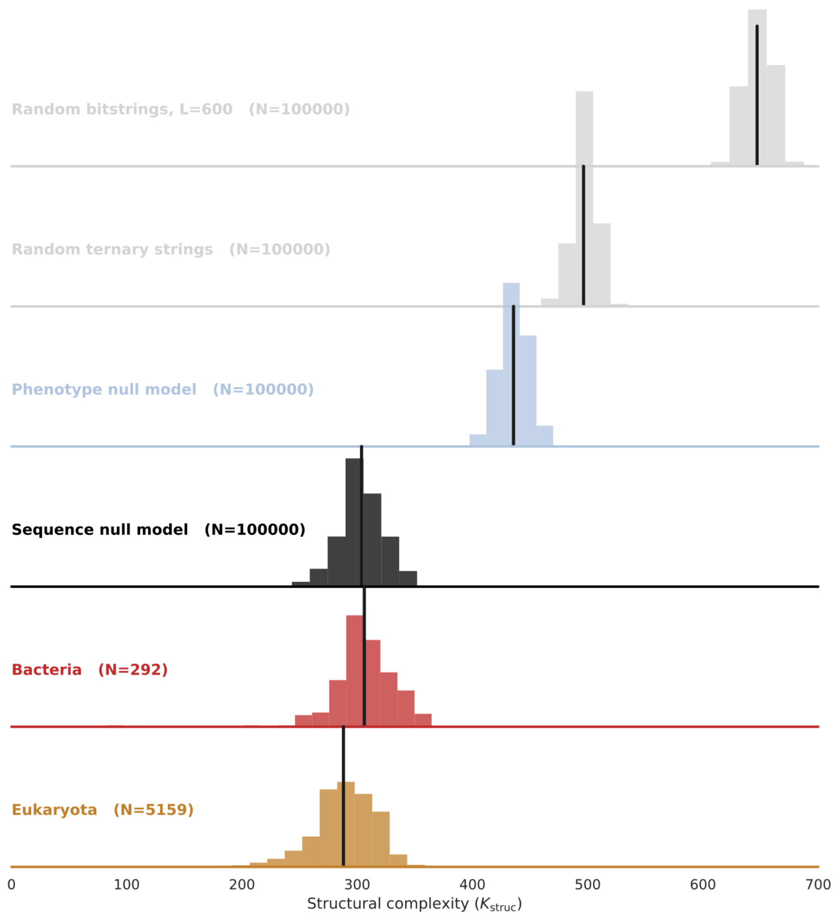


Figure S9 – Probability distribution for structural for the four main RNA sets (same as in Fig. 1–2 in the main text). We also inclue two more abstract sets of strings: Random ternary strings are random strings consisting of '(', ')' and '.'; Random bit strings are random binary strings consisting of '1' and '0' of length $L = 600$, which is the length of the binary strings to which all other five sets are converted before complexity calculation. These comparisons show that from an information theory perspective, the physically possible phenotypes of the random phenotypes sample set are already constrained relative to all possible random strings yielding lower complexity values.

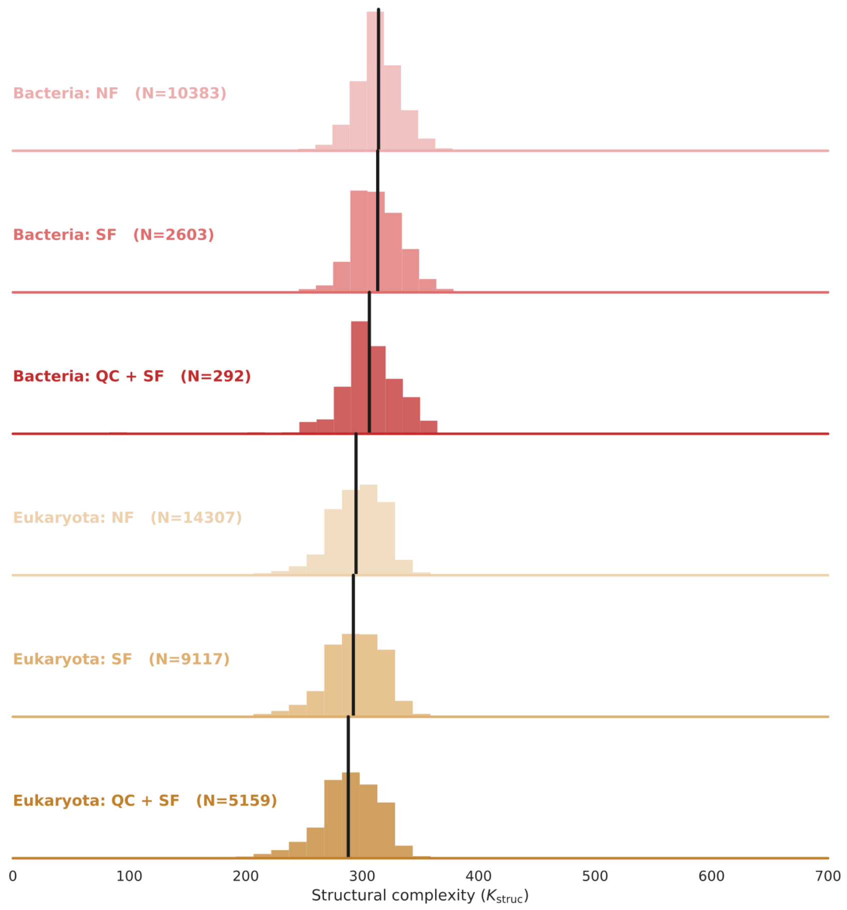


Figure S10 – Effect of filtering on probability distribution of structural complexity. NF: no filtering, QC: quality control, SF: similarity filter, i.e. filter out sequences that have 80% similarity to another sequence in the set. Filtering leads to lower average structural complexity, indicating that low-quality sequences (e.g. partial sequences) and over-sampled sequences are biased towards higher complexity phenotypes.

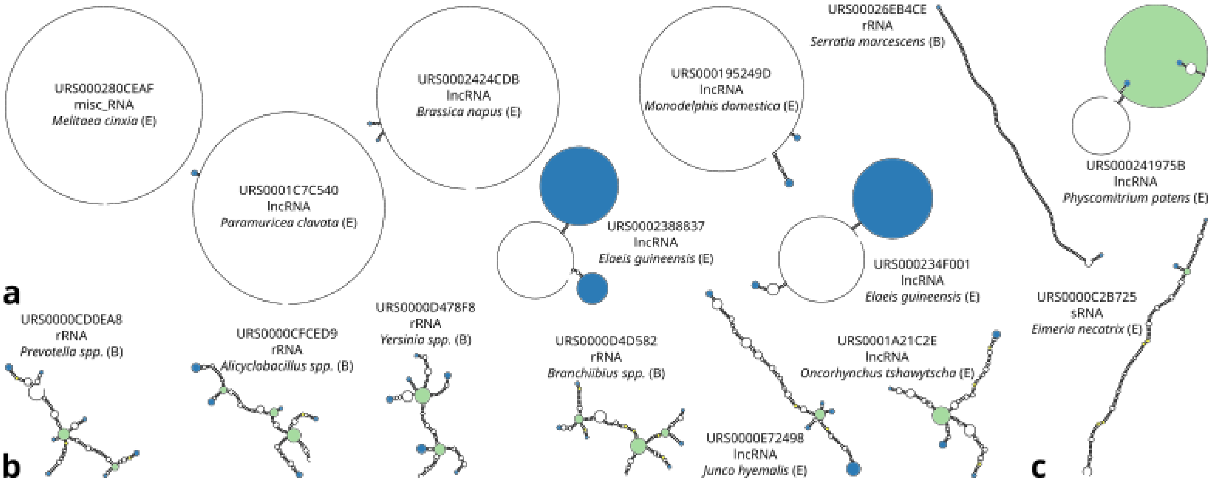


Figure S11 – Example RNA secondary structures of $L = 300$, sequences obtained from RNAcentral, folded using RNAfold (see turqoise markers in Fig. 1c in main text) and visualized using forna, two tools from the ViennaRNA package (Lorenz et al., 2011). (**a**) Eight RNA molecules with the simplest secondary structure, i.e. lowest structural complexity (ordered from low to high). (**b**) Six RNA molecules with the most complex secondary structure, i.e. highest structural complexity (ordered from low to high). (**c**) RNA molecule with the simplest sequence, i.e. lowest sequence complexity. See Table S1 for data. External loops, bulges and junctions are colored as in Fig. 2 in main text.

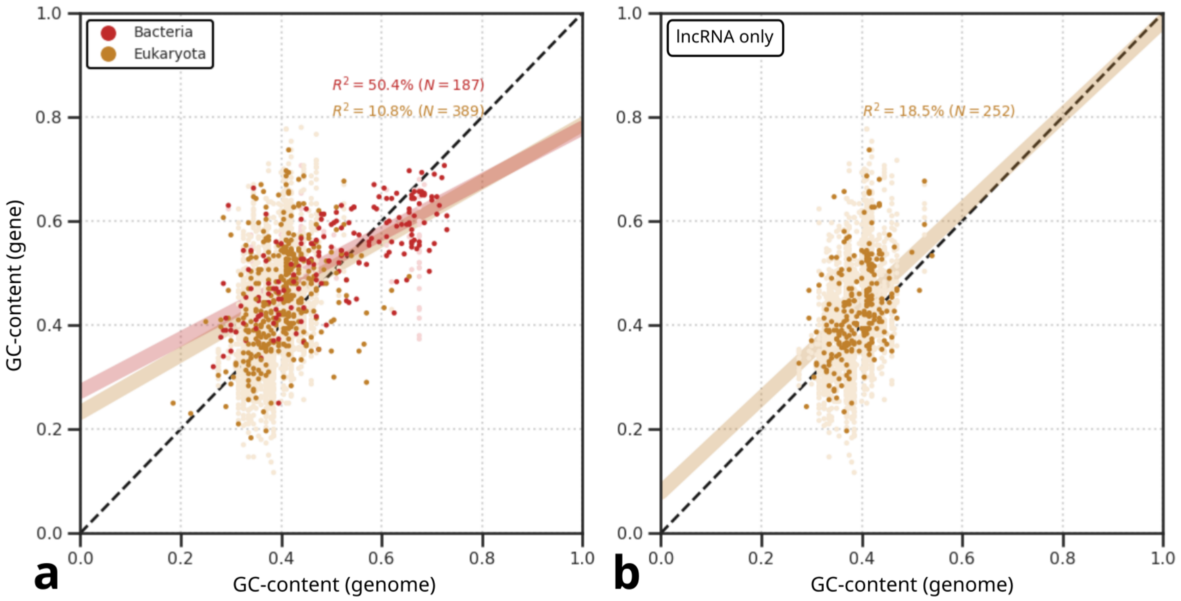


Figure S12 – GC-content of RNA genes correlates with GC-content of the entire genome. (**a**) Across RNA types, the slope of regression lines for bacteria and eukaryota are 0.51 and 0.55, indicating that GC-content of functional RNA genes is only partially driven by genome-scale mutational bias. (**b**) For eukaryotic lncRNA, the slope of the regression line is 0.91, showing that GC-content of lncRNA follows on average the genomic GC-content. However, large variation around the regression line could still point at an impact of selection on GC-content of lncRNA genes. Genomic GC-content was available for 576 taxa in our RNAcentral $L = 300$ dataset. For the fitting, we sampled one RNA gene for each taxon; in the lighter colors in the background all RNA genes for the 576 taxa are shown ($N = 4332$, i.e. accounting for 79% of sequences in our $L = 300$ dataset).

Table S3 – List of annotated archaeal species in our LSU rRNA analysis with very simple structures and those with relatively complex structures (relative structural complexity, $Z_{\mathrm{K}}^{\mathrm{struc}}$). Most archaeal rRNA secondary structures are more robust to mutations ($\lambda$) than an average molecule of similar length ($\lambda \approx 27.5$) and more robust to changes in temperature than the average LSU rRNA ($\langle S \rangle \approx 0.77$). Low structural complexities are the result of low relative sequence complexities ($Z_{\mathrm{K}}^{\mathrm{seq}}$), even in comparison with a shuffled copy which has the same nucleotide bias (i.e. $\Delta Z_{\mathrm{K}}^{\mathrm{seq}}$). Niches were obtained through online literature searches: HTP (hyperthermophile), TP (thermophile), MP (mesophile), TAP (thermoacidophile), U (unknown). Energetic stability is calculated as the energy difference between the minimum-free-energy structure and the average of the ensemble obtained using RNAsubopt by sampling 100 secondary structures from each sequence proportional to free energy. Thermodynamic robustness was calculated as the mean structural similarity of the minimum-free-energy structures at temperatures $T = [27.75, 46.25, 55.5, 64.75, 74]$ with the structure obtained at default temperature ($T = 37$).

| **Species** | $Z_{\mathbf{K}}^{\mathbf{struc}}$ | $Z_{\mathbf{K}}^{\mathbf{seq}}$ | $\Delta Z_{\mathbf{K}}^{\mathbf{seq}}$ | **L** | **GC-content** | $\lambda$ | $\Delta E$ | $\langle S \rangle$ | **Niche** |
|---|---|---|---|---|---|---|---|---|---|
| *Pyrobaculum ferrireducens* | -3.8 | -6.1 | -2.0 | 3027 | 68.1 | 38 | -57.5 | 0.91 | HTP |
| *Staphylothermus marinus* | -3.4 | -7.2 | -2.9 | 3240 | 66.7 | 37 | -60.4 | 0.89 | HTP |
| *Sulfolobus* sp. A20 | -3.3 | -4.2 | -2.3 | 2980 | 61.9 | 39 | -54.3 | 0.90 | HTP |
| *Candidatus* Odinarchaeum yellowstonii | -3.1 | -0.8 | -1.1 | 3073 | 55.9 | 28 | -63.4 | 0.85 | TP |
| *Pyrobaculum aerophilum* | -3.1 | -6.6 | -2.4 | 3022 | 67.9 | 35 | -55.7 | 0.87 | HTP |
| *Caldivirga* sp. MG_3 | -2.9 | -3.6 | -3.7 | 3353 | 60.2 | 22 | -76.6 | 0.85 | HTP |
| *Vulcanisaeta* sp. AZ3 | -2.7 | -7.7 | -5.5 | 2916 | 64.6 | 32 | -57.2 | 0.89 | HTP |
| *Desulfurococcus amylolyticus* | -2.7 | -5.4 | -1.6 | 3093 | 66.0 | 38 | -56.2 | 0.87 | HTP |
| *Methanopyrus* sp. KOL6 | -2.7 | -8.2 | -4.6 | 3092 | 68.8 | 37 | -51.6 | 0.89 | HTP |
| *Desulfurococcus amylolyticus* | -2.6 | -5.7 | -0.5 | 3085 | 65.8 | 37 | -58.2 | 0.89 | HTP |
| *Methanopyrus* sp. SNP6 | -2.5 | -8.4 | -3.9 | 3092 | 68.8 | 30 | -50.3 | 0.90 | HTP |
| *Methanothrix* | 0.5 | 0.0 | 0.5 | 2862 | 53.2 | 40 | -67.0 | 0.80 | TP |
| *Candidatus* Altarchaeales | 0.5 | -1.0 | -1.1 | 2629 | 47.1 | 38 | -68.3 | 0.80 | U |
| *Methanothermobacter* sp. CaT2 | 0.5 | -3.6 | -2.1 | 3017 | 56.5 | 31 | -70.3 | 0.79 | TP |
| *Methanolobus* sp. T82-4 | 0.7 | -0.3 | -0.5 | 2914 | 51.5 | 43 | -67.7 | 0.81 | U |
| *Ferroplasmaceae* | 0.7 | -0.4 | -0.4 | 2897 | 47.4 | 27 | -76.0 | 0.81 | MP |
| *Methanobacterium* sp. BRmetb2 | 0.7 | -1.3 | -1.6 | 2847 | 51.1 | 38 | -66.5 | 0.84 | U/MP |
| *Candidatus* Methanoperedens nitroreducens | 1.0 | -0.7 | 0.4 | 2894 | 53.6 | 30 | -65.8 | 0.85 | MP |
| *Methanothrix harundinacea* | 1.1 | 0.0 | 0.4 | 2764 | 54.3 | 31 | -65.1 | 0.81 | MP |
| *Metallosphaera sedula* | 1.2 | -3.6 | -2.1 | 2903 | 61.1 | 32 | -58.1 | 0.82 | TP |
| *Thermoplasma acidophilum* | 2.4 | -1.5 | -1.2 | 2913 | 53.0 | 27 | -69.0 | 0.82 | TAP |

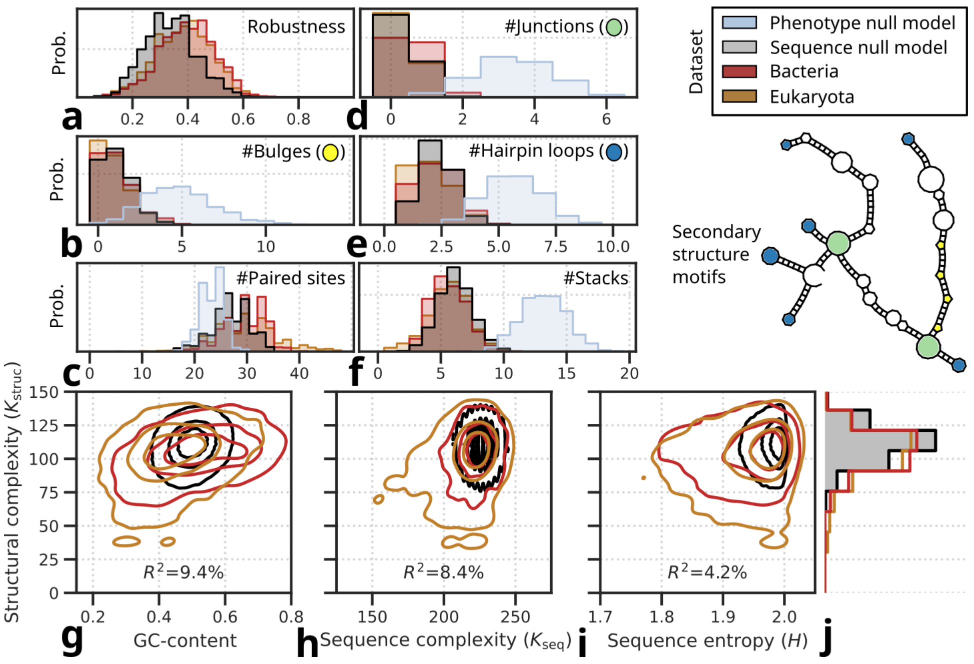


Figure S13 – Characterization of mutational robustness, structural motifs and sequence features in random and natural RNA of length $L = 100$, for comparison with analysis of RNA of length $L = 300$ in the main text (Fig. 2). (**a**) Probability distribution of mutational robustness (fraction of mutants with identical secondary structure) for the three sequence sets. (**b**–**f**) Probability distributions of secondary structure motif counts. Note the effect of developmental bias, defined as the difference between the phenotype null-model and the sequence null-model. (**g**–**j**) Kernel density plots showing correlations of structural complexity with three features of sequence composition, i.e. (**g**) GC-content, (**h**) sequence complexity, and (**i**) sequence entropy. Together, these features account for 22% of the total variation in structural complexity; similarly, $R^2$ values represent variation explained across all three RNA sequence sets.

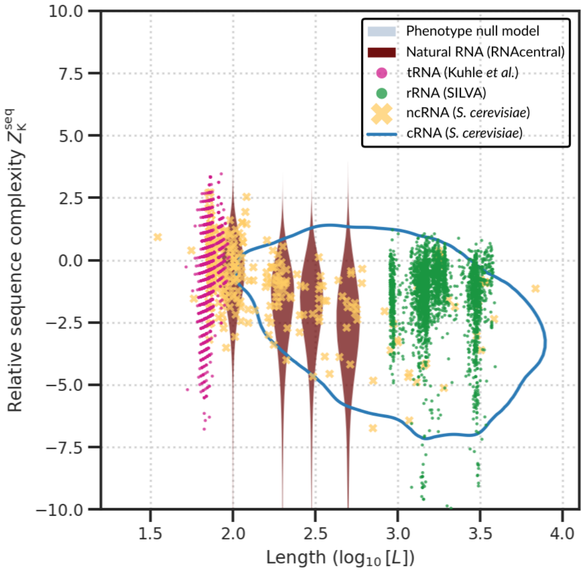


Figure S14 – Scaling of sequence complexity with length (similar to scaling of structure complexity with length; Fig. 3 in main text). Except for random phenotypes—which do not have a corresponding sequence and are thus absent here—and the smallest random genotype set ($L = 10$)—for which most sequences form no base pairs in their structure and are thus removed from the fitting of structural complexity with length (Fig. 3)—the plot shows the same RNA molecules as in Fig. 3.

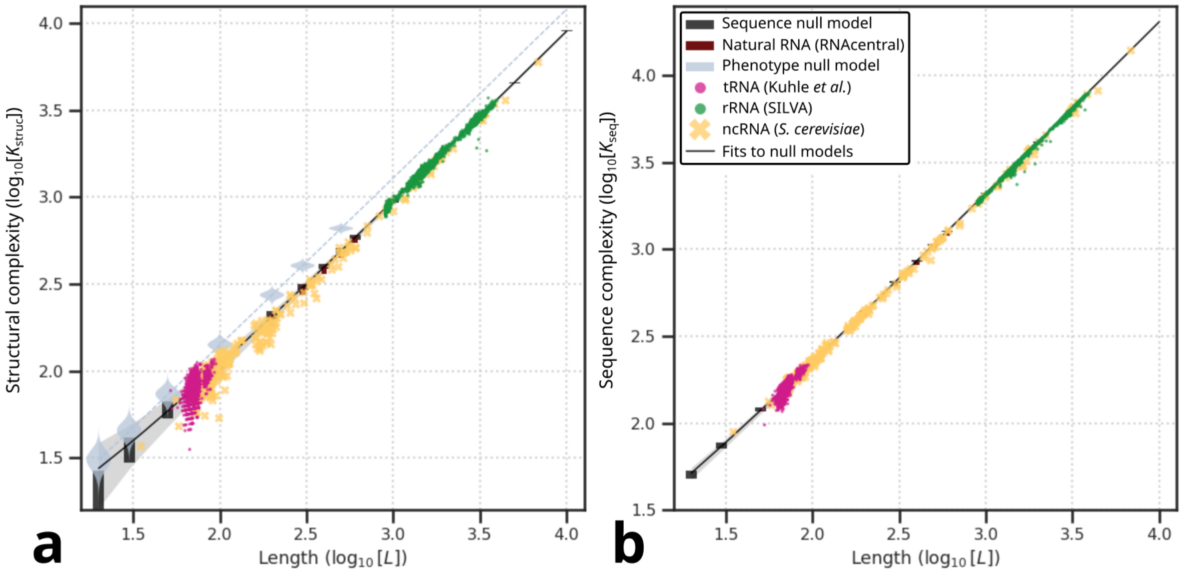


Figure S15 – Structural complexity and sequence complexity scale with sequence length. Fig. 3, S14 show structural complexity and sequence complexity as normalized by the power law fit for random genotype samples at different specific sequence lengths. See Fig. S16 for the goodness of fit.

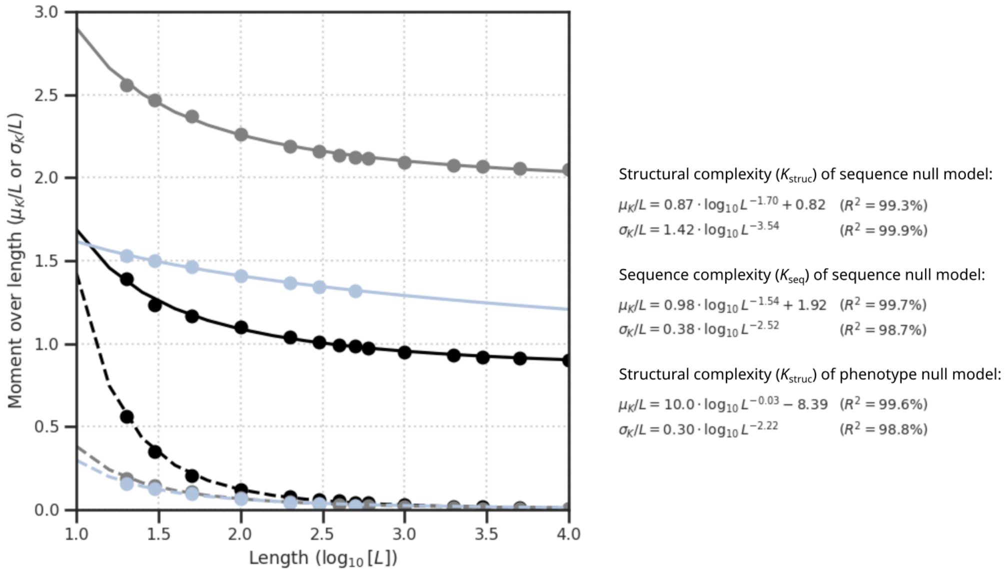


Figure S16 – Fitting power law relationship for structural and sequence complexity against length using the `curve_fit` function from the scipy python package (Virtanen et al., 2020). Means and standard deviations of structural and sequence complexity (see Fig. S3) are fitted to the functional form $f(x) = a \cdot \log_{10} L^b + c$, fixing $c = 0$ for standard deviations to keep them strictly positive. Line colors denote distributions being fit (black: structural complexity of random genotypes, grey: sequence complexity of random genotypes, light blue: structural complexity of random phenotypes). Line types denote moments being fit (solid: mean, striped: standard deviation). Note that we have random genotype samples of lengths ranging four orders of magnitude; random phenotype samples range almost three orders of magnitude in length.

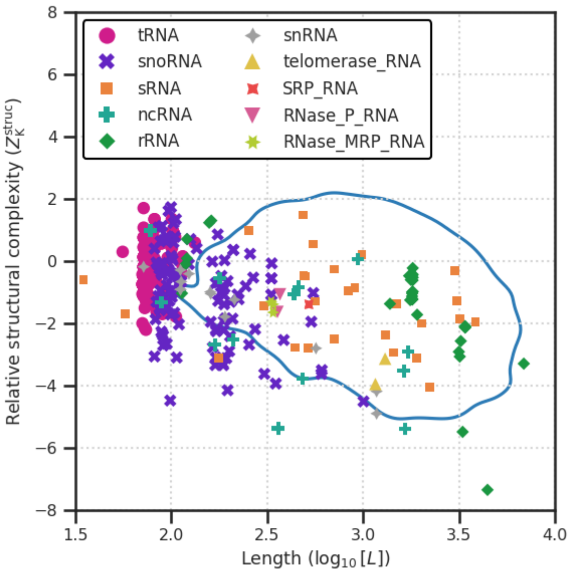


Figure S17 – RNA types of *S. cerevisiae* differ in sequence length and relative structural complexity. Blue line shows the contour for coding (i.e. non-functional) RNA. Data represent all genes (347 non-coding RNA and 5521 coding RNA) of the *S. cerevisiae* genome, taken from SGD (Engel et al., 2022).

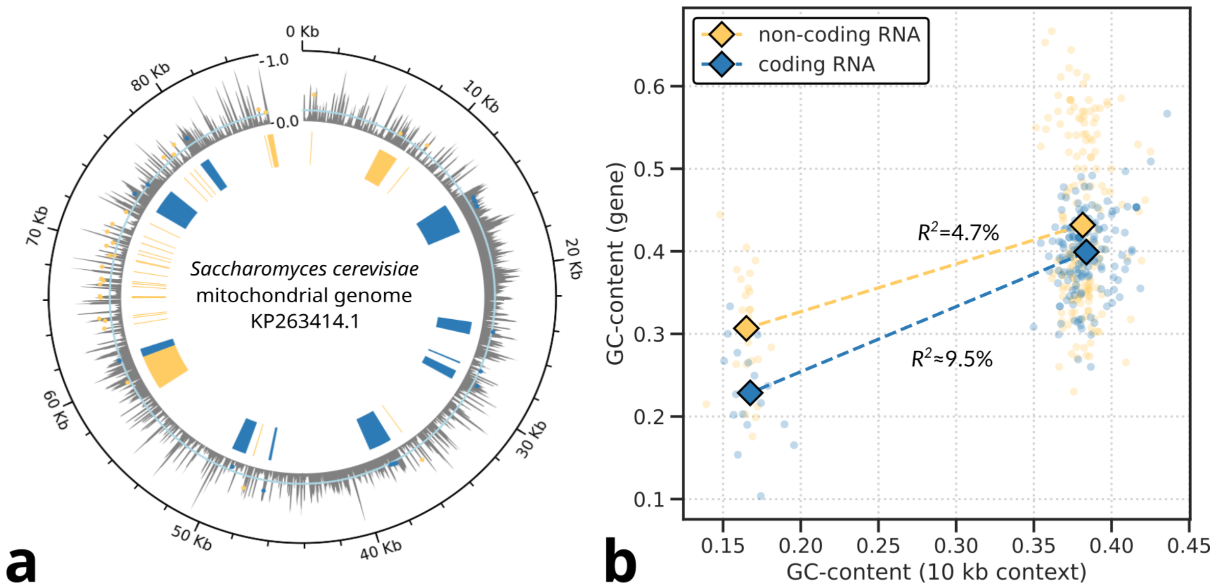


Figure S18 – Non-coding RNA genes from the mitochondrial genome of yeast have elevated GC-content, exposing selection for structural complexity. The low GC-content of the mitochondrial genome yields low structural complexity of coding RNA genes, but non-coding RNA genes maintain sufficiently high GC-content to conserve structural complexity to similar levels as in the nuclear genome (cf. Fig. 4). (**a**) Distribution of genes on mitochondrial genome, with variation in GC-content (grey shows mean GC-content per 100-nt frame; blue denotes genomic average).(**b**) Correlation between GC-content of genes and that of their direct genomic context (10kb windows around the focal gene) for mitochondrial and nuclear genes. R-squared indicates variance explained by Spearman rank correlation. For coding RNA, the mean R-square of 100 bootstrapped samples is shown.

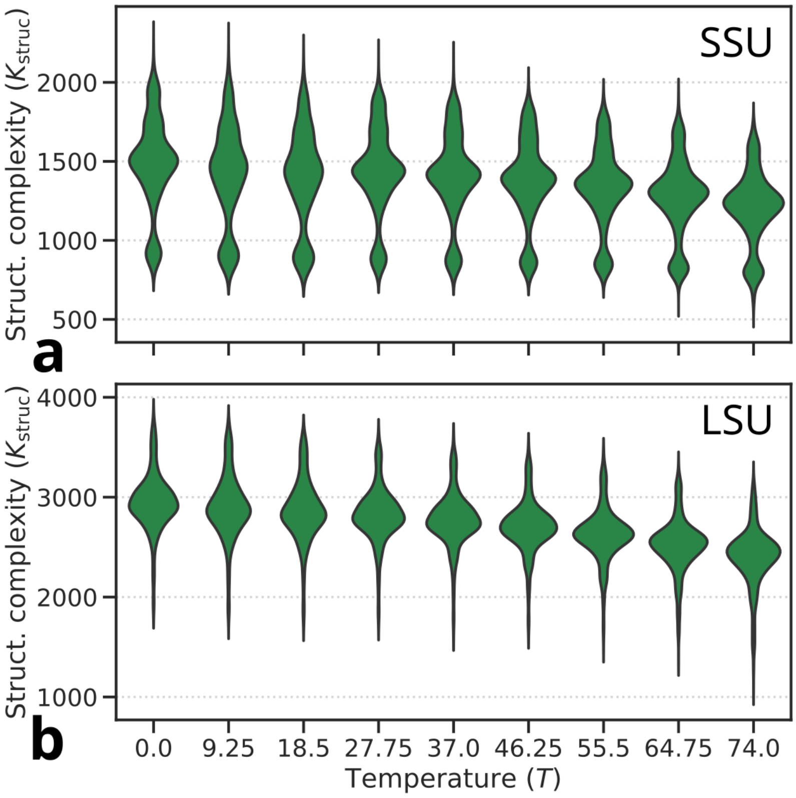


Figure S19 – Impact of temperature on RNA folding: with increasing temperature the minimum-folding-energy structures of rRNA sequences become simpler. Default temperature, used throughout this work, is at $T = 37$.

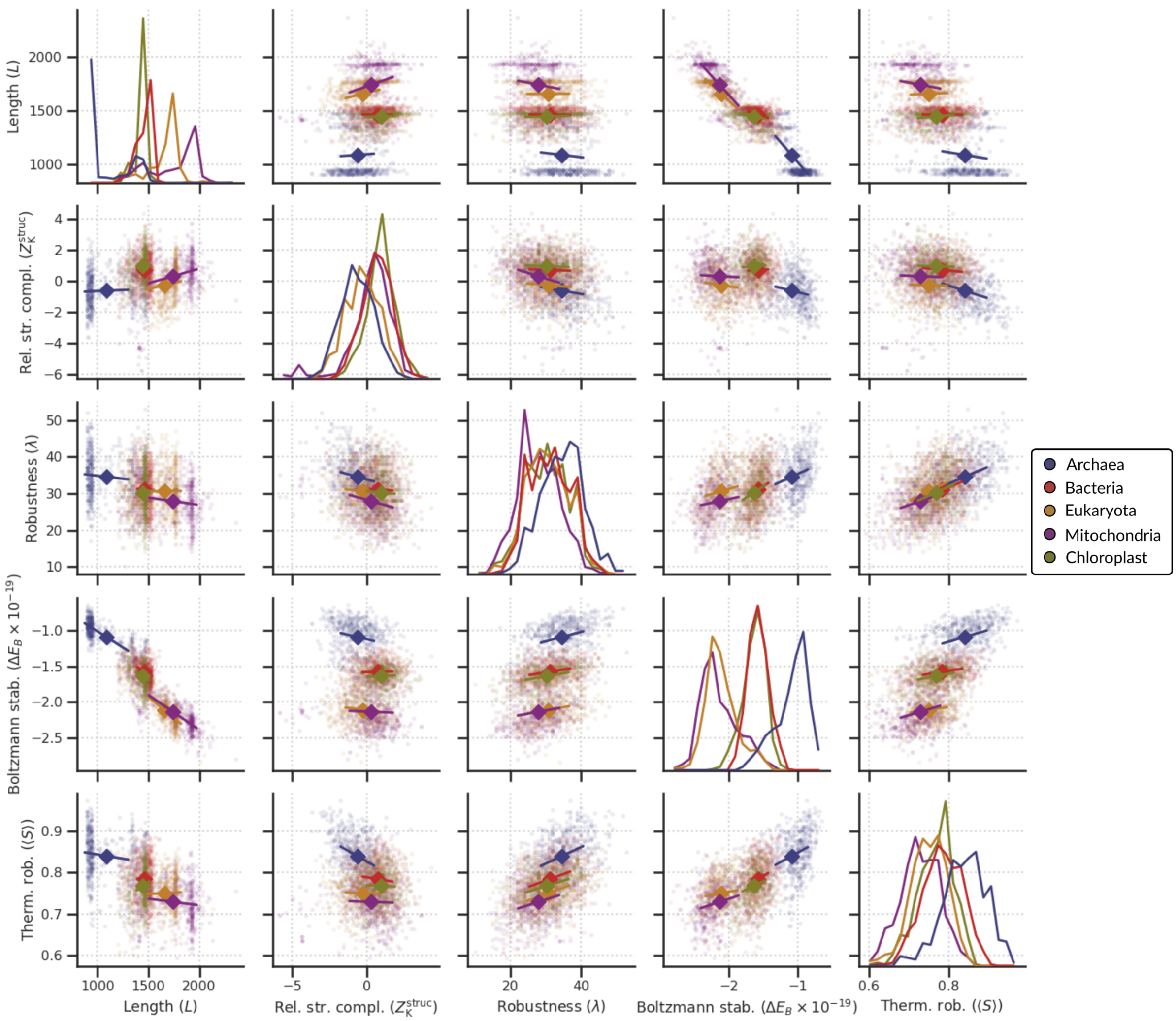


Figure S20 – Correlations between complexity, robustness and stability in SSU rRNA. Length, relative structural complexity and mutational robustness are as described in the main text (e.g. Fig. 5). Energetic stability is calculated as the energy difference between the minimum-free-energy structure and the average of the ensemble obtained using RNAsubopt by sampling 100 secondary structures from each sequence proportional to free energy. Thermodynamic robustness was calculated as the mean structural similarity of the minimum-free-energy structures at temperatures $T = [27.75, 46.25, 55.5, 64.75, 74]$ with the structure obtained at default temperature ($T = 37$).

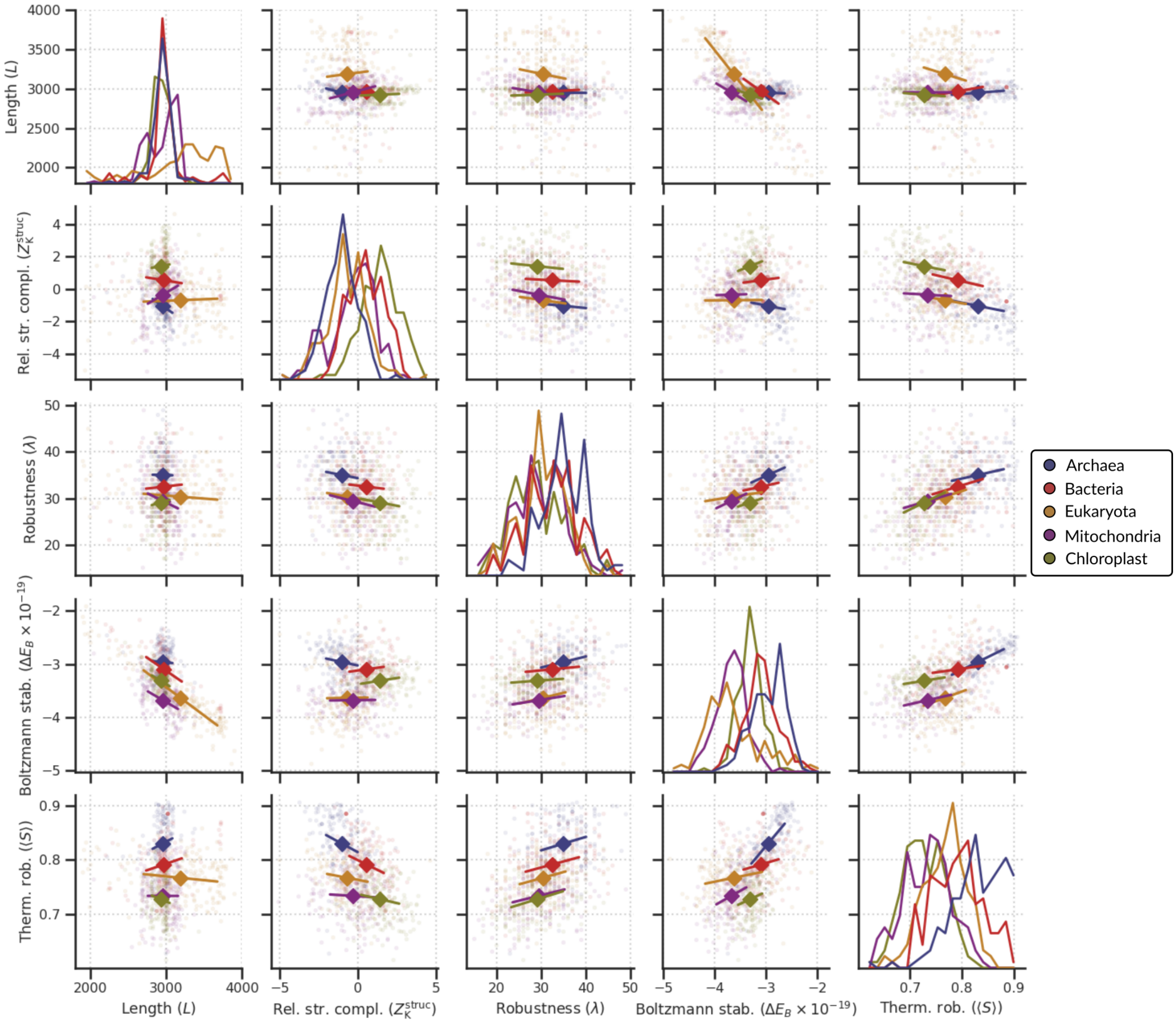


Figure S21 – Correlations between complexity, robustness and stability in LSU rRNA. Length, relative structural complexity and mutational robustness are as described in the main text (e.g. Fig. 5). Energetic stability is calculated as the energy difference between the minimum-free-energy structure and the average of the ensemble obtained using RNAsubopt by sampling 100 secondary structures from each sequence proportional to free energy. Thermodynamic robustness was calculated as the mean structural similarity of the minimum-free-energy structures at temperatures $T = [27.75, 46.25, 55.5, 64.75, 74]$ with the structure obtained at default temperature ($T = 37$).

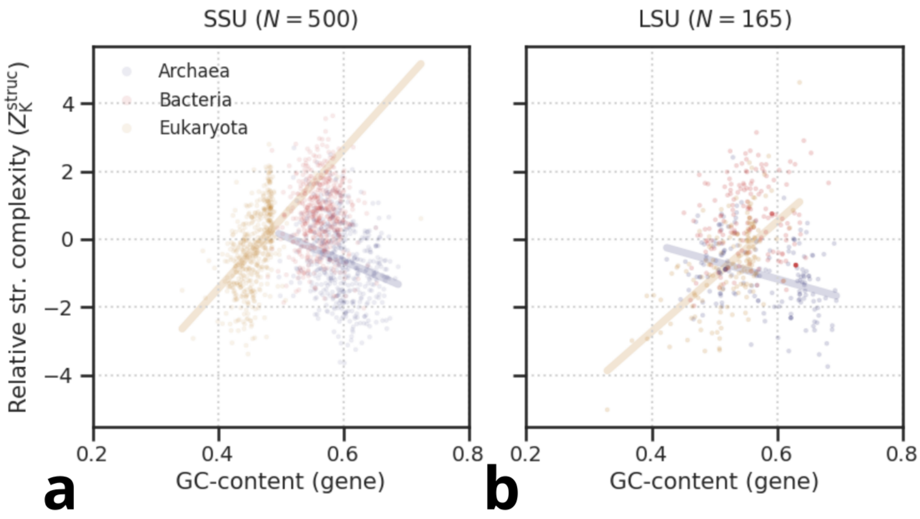


Figure S22 – Structural complexity scales non-monotonically with GC-content in rRNA if archaea with high GC-content are included. Linear fits are plotted through subsets if there is a significant Spearman rank correlation ($p < 0.05$).
